# Conformational surveillance of Orai1 by a rhomboid intramembrane protease prevents inappropriate CRAC channel activation

**DOI:** 10.1101/2020.04.04.025262

**Authors:** Adam G Grieve, Yi-Chun Yeh, Lucrezia Zarcone, Johannes Breuning, Nicholas Johnson, Kvido Stříšovský, Marion H Brown, Anant B Parekh, Matthew Freeman

**Affiliations:** Sir William Dunn School of Pathology, University of Oxford, UK; Department of Physiology, Anatomy and Genetics, University of Oxford, UK; Institute of Organic Chemistry and Biochemistry of the Czech Academy of Sciences (IOCB), Czech Republic; GlaxoSmithKline (GSK), UK; Cancer Research UK Beatson Institute, UK

**Keywords:** Intramembrane protease, rhomboid, RHBDL2, calcium signalling, Orai1, Stim1, CRAC channel, membrane protein, homeostasis, T cells

## Abstract

Calcium influx through plasma membrane calcium release-activated calcium (CRAC) channels, which are formed of hexamers of Orai1, is a potent trigger for many important biological processes, most notably in T cell mediated immunity. Through a bioinformatics-led cell biological screen, we have identified Orai1 as a substrate for the rhomboid intramembrane protease, RHBDL2. We show that RHBDL2 prevents stochastic signalling in unstimulated cells through conformational surveillance and cleavage of inappropriately activated Orai1. A conserved, disease-linked proline residue is responsible for RHBDL2 recognising only the active conformation of Orai1, and cleavage by RHBDL2 is required to sharpen switch-like signalling triggered by store-operated calcium entry. Loss of RHBDL2 control of Orai1 causes severe dysregulation of CRAC channel effectors including transcription factor activation, inflammatory cytokine expression and T cell activation. We propose that this seek-and-destroy function may represent an ancient activity of rhomboid proteases in degrading unwanted signalling proteins.

## Introduction

Signalling controls most cellular functions and must therefore be precisely regulated in time and space. Although some signals produce graded responses, most are converted into binary outputs: having received an input, usually a ligand binding to a receptor, a cell changes its state in a switch-like way. Much signalling is triggered by integral membrane receptors including, for example, growth factor and cytokine receptors, ion channels, and G protein coupled receptors. To maintain switch-like signalling, it is essential that spontaneous activation is prevented in the absence of stimulus, but the post-translational control mechanisms for surveillance and prevention of inappropriate signalling are unknown. More generally, little is known about the regulation of the abundance and activity of most cell surface signalling proteins.

A good example of switch-like signalling is the control of calcium ion (Ca^2+^) flux across the eukaryotic plasma membrane (PM), which acts as a barrier between high extracellular and low cytoplasmic Ca^2+^ concentrations. Almost all cells in the animal kingdom regulate the level of cytosolic Ca^2+^ to control their function: a sharp rise in cytosolic Ca^2+^ controls enzymatic activity, protein-protein interactions, gene activation, cell proliferation and apoptosis (Berridge et al., 2000). One of the primary routes of regulating Ca^2+^ entry in non-excitable cells is via Ca^2+^ release-activated Ca^2+^ (CRAC) channels (Parekh and Putney, 2005). Opening of CRAC channels at the cell surface causes a rapid increase of cytosolic Ca^2+^, which activates many important signalling pathways, the most studied being in the adaptive immune system. The pore-forming subunits of CRAC channels are Orai proteins (Feske et al., 2006; Prakriya et al., 2006; Vig et al., 2006; Zhang et al., 2006). Orai1, the founding and most ubiquitously expressed member of the family, was originally identified as a genetic cause of severe combined immunodeficiency in humans, and plays essential roles in T cell immunity (Feske et al., 2006).

CRAC channel activity must be transient and tightly controlled to prevent aberrant signalling. Indeed, leaky or defective CRAC channel activity is a direct cause of a group of diseases collectively referred to as channelopathies (Feske, 2010). CRAC channels are gated by the store-operated Ca^2+^ entry signalling pathway, in response to the stimulated release of stored endoplasmic reticulum (ER) Ca^2+^ by phospholipase C signalling (Hogan and Rao, 2015; Prakriya and Lewis, 2015). The consequent reduction of ER Ca^2+^ is sensed by the ER-resident membrane protein Stim1 (Hogan and Rao, 2015), resulting in its interaction with Orai1 at PM-ER contact sites, CRAC channel opening, and the influx of extracellular Ca^2+^. Channel opening relies both on Orai1 multimerisation into a pore-forming hexameric unit, and a concerted set of conformational changes (Hou et al., 2012; Cai et al., 2016; Hou et al., 2018). The main interaction with the cytoplasmic domain of Stim1 occurs via the C-terminal cytoplasmic domain of Orai1, which is anchored to the membrane by its fourth transmembrane domain (TMD) (Park et al., 2009). This interaction triggers allosteric activation and opening of CRAC channels (Yeung et al., 2019). Importantly, the correct stoichiometry between Orai1 and Stim1 is essential for normal store-operated Ca^2+^ entry (Mercer et al., 2006; Peinelt et al., 2006; Soboloff et al., 2006; Scrimgeour et al., 2009; Hoover and Lewis, 2011; Yeh et al., 2019). Overall, Stim1 binding and trapping of Orai1 at PM-ER contact sites is the rate-limiting step in CRAC channel activation and is therefore a major regulatory switch for Ca^2+^ influx and downstream signalling.

One class of enzymes that has the capacity to be involved in regulating the signalling function of integral membrane proteins by inactivation or degradation are the intramembrane proteases, which use their active sites in the lipid bilayers of cell membranes to cleave TMDs of substrates. Most known functions of intramembrane proteases are to release signalling domains from membrane-tethered precursors, thereby triggering a signalling event. However, a wider range of roles is becoming apparent, including participating in some forms of ER-associated degradation (Fleig et al., 2012). Rhomboids are evolutionarily widespread intramembrane serine proteases. Despite extensive study and well-understood functions in several species (Freeman, 2014), and some scattered knowledge of their mammalian function (Lohi et al., 2004; Adrain et al., 2011; Fleig et al., 2012; Johnson et al., 2017) a comprehensive understanding of their physiological importance in mammals has been hampered by a lack of validated substrates (Lastun et al., 2016). To date, most identified rhomboid substrates are type I, single-pass transmembrane proteins. One feature that appears to be common to many intramembrane protease substrates is helical instability in transmembrane segments (Ye et al., 2000; Lemberg and Martoglio, 2002; Urban and Freeman, 2003), often created by the presence of helix-breaking residues such as prolines or glycines. The importance of this feature to rhomboid recognition is highlighted not only by their conservation and functional necessity, but also by the observation that otherwise uncleavable TMDs can be converted into rhomboid substrates by the introduction of a proline residue (Moin and Urban, 2012). This raises the possibility that rhomboids may have originally evolved to recognise non-canonical transmembrane helices.

Our overall goal is to discover the conceptual and mechanistic themes associated with rhomboid intramembrane proteolysis and to uncover their physiological roles in mammals. Here, through a bioinformatics-led cell biological screen, we identify the fourth TMD of Orai1 as a substrate for the PM localised rhomboid protease RHBDL2. We show that proteolysis of Orai1 by RHBDL2 sharpens the precision of store-operated Ca^2+^ entry by preventing stimulus-independent CRAC channel activation and inflammatory cytokine expression in unstimulated cells. Mechanistically, RHBDL2 prevents this inappropriate signalling by conformational selection of the activated form of Orai1. The pathophysiological importance of this mechanism is highlighted by our demonstration that an activating disease-associated proline-to-leucine mutation in Orai1 TMD4 prevents RHBDL2 recognition, and severe defects in primary T cell activation occur upon RHBDL2 loss. We propose that conformational surveillance of polytopic proteins may represent an ancient rhomboid protease activity that could predate its better-known roles in the cleavage and extracellular release of signalling molecules.

## Results

### Orai1 is an RHBDL2 substrate

To discover new substrates for the rhomboid intramembrane protease, RHBDL2, we used a bioinformatic approach, followed by functional validation with a cell-based assay. We focused on three characteristics of known rhomboid substrates: their TMDs have a type I orientation (NH^2^-out, COOH-in), many contain extracellular EGF-like domains and, like substrates of intramembrane proteases in general, they often contain helix-destabilising amino acids (Freeman, 2014). Therefore, we identified candidates by the presence of extracellular EGF domains and/or through profile-profile alignments with the online server HHpred (Zimmermann et al., 2018) to find type I TMDs that have structural similarity to one of the best characterised rhomboid substrates, *Drosophila melanogaster* Spitz (Freeman, 2014).

The top bioinformatic TMD hits (approximately 175, **Table S1**) were inserted into a reporter that was co-expressed in cells with RHBDL2, so that RHBDL2-dependent cleavage leads to accumulation of extracellular alkaline phosphatase (AP) (**Figure 1A;** *left*), which can be detected using a colorimetric phosphatase assay. As a positive control, we used the TMD of Spitz, which can be cleaved by RHBDL2 (**Figure 1B, Figure S1A**) (Urban and Freeman, 2003; Strisovsky et al., 2009). Among the strongest validated hits, we found the fourth TMD of all three members of the Orai family of Ca^2+^ channels (**Figure 1A, 1C**). The cleavage required the catalytic serine residue of RHBDL2, as well as the hallmark rhomboid WR motif in the L1 loop that connects TMD1 and TMD2 (**Figure 1D**) (Lemberg and Freeman, 2007).

**Figure 1:**
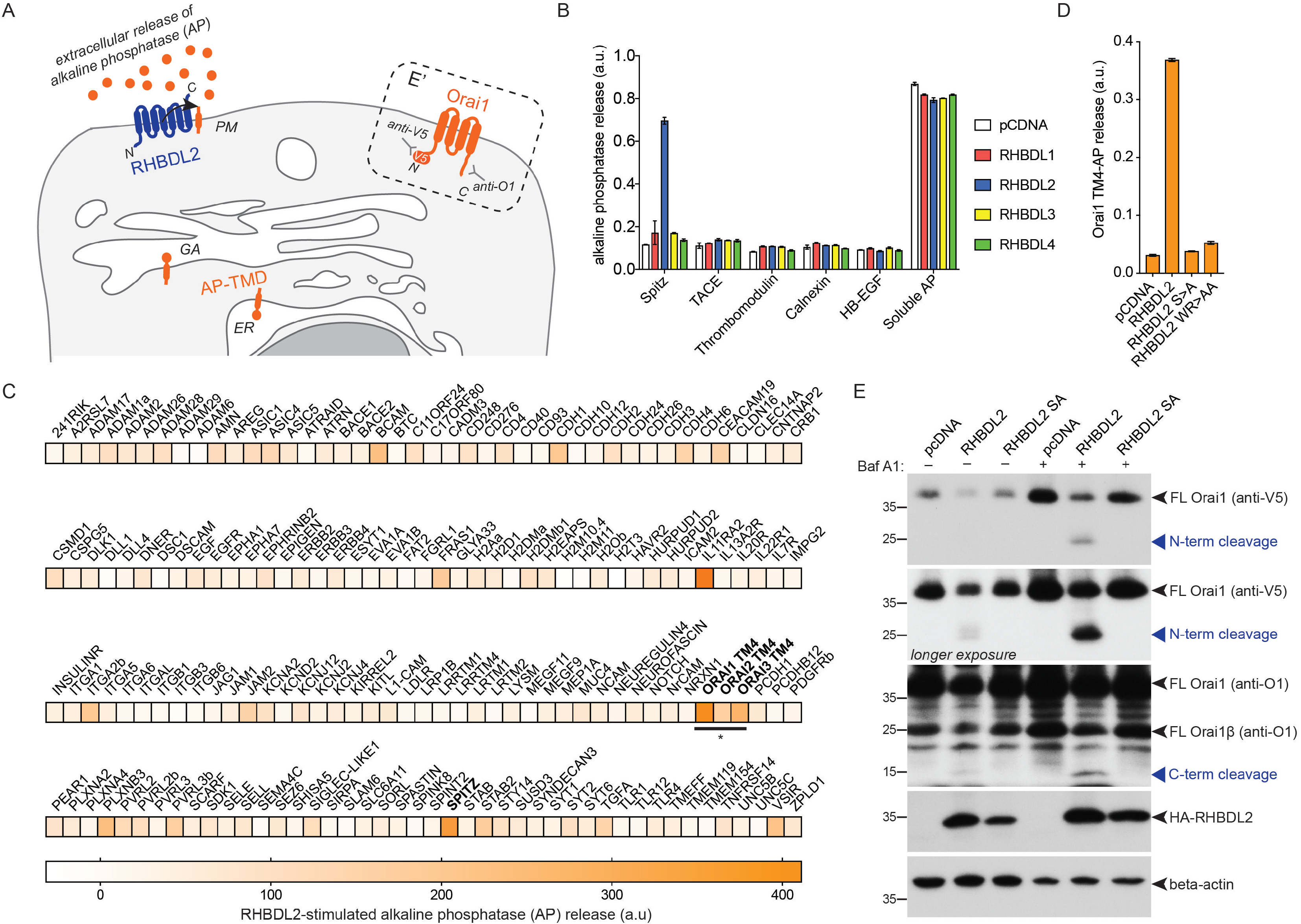
Orai1 is an RHBDL2 substrate. **A**. Scheme of the alkaline phosphatase-transmembrane domain (AP-TMD) screen. Each AP-TMD (orange) has a signal sequence ensuring their insertion within the endoplasmic reticulum (ER) and subsequent delivery to the cell surface. Upon co-expression with RHBDL2 (blue), if AP-TMD is cleaved, it will release AP into the extracellular medium. In the dashed box (E’), the topology of V5-Orai1 is illustrated, indicating the epitopes recognised by antibodies used for western blot in panel E. GA = Golgi apparatus. PM = plasma membrane. **B-D.** HEK293 cells were transfected with pcDNA, 3xHA-RHBDL1-4 or RHBDL2 mutants (S->A, or WR->AA) and indicated AP-TMDs for 48 hours. Soluble AP is AP with a signal sequence, but no transmembrane anchor. Released AP was collected over the final 16 hours of expression. Values represent the level of released AP/total AP (**B, D**) or in (**C**) these values were converted into a normalised level of “RHBDL2-stimulated AP release” (for each AP-TMD: AP release upon RHBDL2 expression was divided by values taken for pCDNA transfected controls, and multiplied by 100. All values have AP release with pCDNA subtracted, as this release is RHBDL2-independent). For the AP-TMD screen, n = 2 biological repeats for each AP-TMD. Error bars indicate standard error of the mean. **E**. Western blots of lysates from HEK293 cells transfected with V5-Orai1 and indicated RHBDL2 constructs for 48 hours, treated with 100 nM Bafilomycin A1 for 16 hours, and probed with Orai1, V5, HA or beta-actin antibodies. Different full-length forms of Orai1 (Orai1β arises from alternative start sites methionine-64 or -71 (Fukushima et al., 2012)) and their cleavage products are indicated by the blue arrowheads.

We tested whether RHBDL2 could cleave full length Orai1, which unlike most known rhomboid substrates is a polytopic protein (**Figure 1A;** *E’*). Upon expression with RHBDL2, full length Orai1 and its short isoform (Orai1β) were cleaved into two fragments of molecular weights that confirmed cleavage within the fourth TMD (**Figure 1E**). Immunofluorescent labelling showed that cleavage led to Orai1 internalisation from the PM, suggesting that it was degraded as a consequence of cleavage (**Figure S1B**). Accordingly, treatment of cells with Bafilomycin A1 – a lysosomal degradation inhibitor – increased the level of the N- and C-terminal Orai1 cleavage products (**Figure 1E**). Rhomboid substrates are normally cleaved between amino acids with small side chains, and bulky residues at the cleavage site often render them uncleavable (Urban and Freeman, 2003). HHpred-generated alignments between TMD4 of Orai1 and the Spitz TMD showed a perfect alignment of the alanine-serine cleavage site in Spitz with alanine-238/serine-239 in Orai1 (Strisovsky et al., 2009) **(Figure S1C)**. Consistent with the expectation that they comprise the site of Orai1 cleavage, both A238F and S239F mutations blocked RHBDL2-dependent proteolysis of Orai1 TMD4 (**Figure S1D**). Overall, these results confirm that the fourth TMD of Orai1 is a *bona fide* substrate of RHBDL2, and that cleavage triggers subsequent Orai1 degradation in lysosomes.

### RHBDL2 controls CRAC channel activity

The fourth TMD of Orai1 both anchors the cytoplasmic C-terminal Stim-interacting domain and is proposed to be central to the conformational changes that initiate CRAC channel opening (Park et al., 2009; Yeung et al., 2019) (**Figure 2A**). We therefore tested the effect of RHBDL2 expression on endogenous store-operated Ca^2+^ entry by monitoring cytosolic Ca^2+^ with the reporter dye Fura-2. Signalling was triggered by thapsigargin treatment, which depletes ER Ca^2+^, followed by addition of physiological levels of extracellular Ca^2+^. We found that RHBDL2 expression reduced store-operated Ca^2+^ entry by ~50%, a level similar to that observed upon Orai1 depletion by siRNA (**Figure 2B-2D**). In these standard assays, cytosolic Ca^2+^ reflects a balance of Ca^2+^ influx and efflux, thus making it formally possible that the observed difference was not the direct result of altered CRAC channel activity, but instead accelerated Ca^2+^ efflux by PM Ca^2+^-ATPases. To discriminate between these two scenarios, we assayed influx of barium ions (Ba^2+^), which can be transported through CRAC channels but cannot be pumped out of the cell by PM Ca^2+^-ATPases (Hoth, 1995; Bakowski and Parekh, 2007). Again, there was a ~50% diminished influx of Ba^2+^ after thapsigargin treatment, supporting the conclusion that RHBDL2 expression prevented CRAC channel activity directly (**Figure 2E, 2F**). To investigate the functional correlate of this effect, we examined the regulated translocation of the transcription factor NFAT (nuclear factor of activated T cells), which is triggered by CRAC channel activity (Hogan et al., 2010; Kar et al., 2011; Kar and Parekh, 2013). Expression of RHBDL2 – but not the related rhomboid protease RHBDL4 – inhibited GFP-NFAT nuclear translocation upon treatment with thapsigargin (from ~98% nuclear NFAT upon thapsigargin treatment in control to ~43% in RHBDL2 expressing cells) (**Figure 2G**). We conclude that RHBDL2 cleavage of Orai1 prevents both endogenous CRAC channel activity and the stimulated nuclear translocation of NFAT.

**Figure 2:**
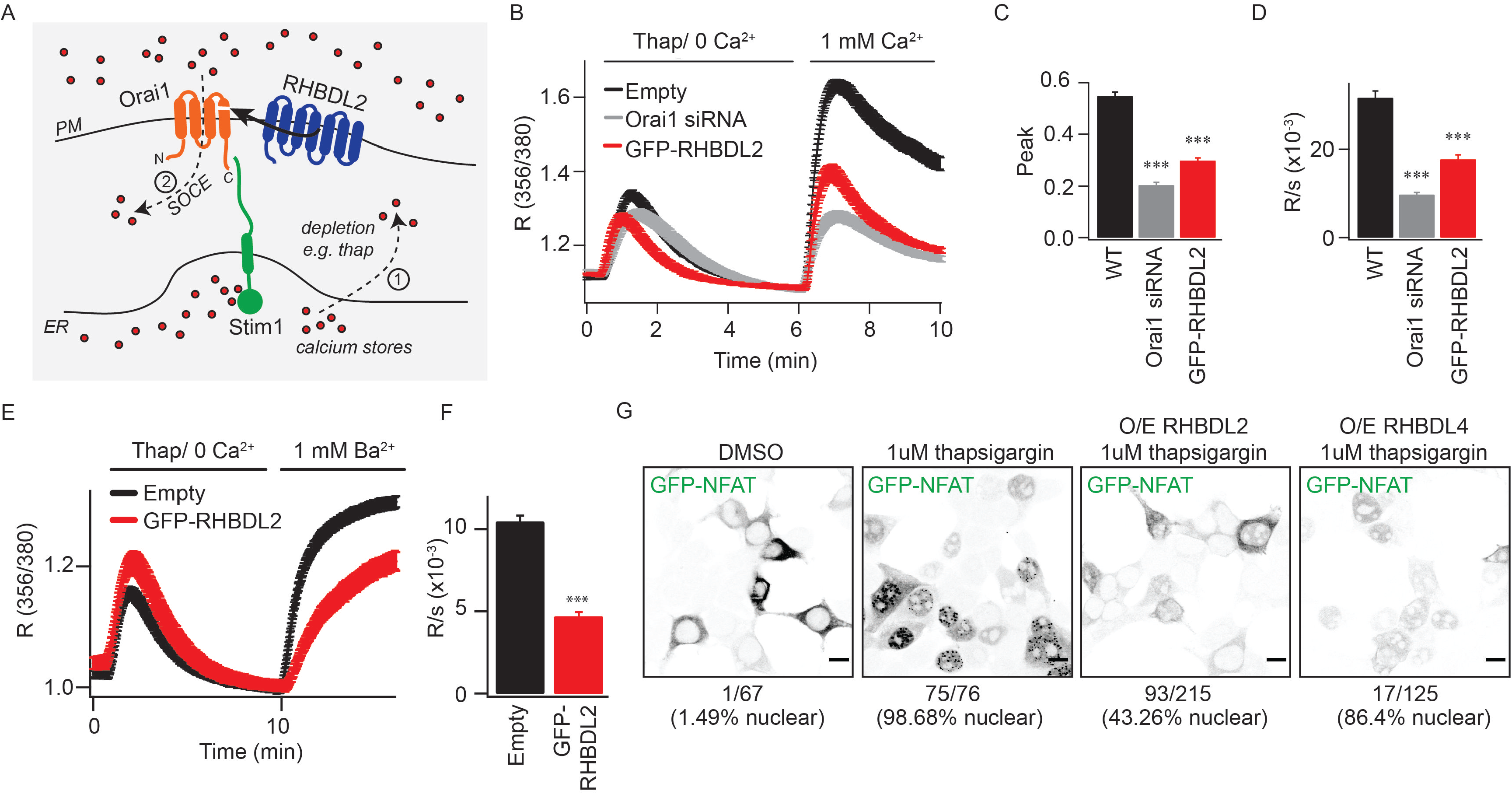
RHBDL2 downregulates CRAC channel activity. **A.** Overview of the store-operated Ca^2+^ entry pathway. Upon depletion of ER Ca^2+^ stores (red dots) by thapsigargin (step 1), Stim1 (green) oligomerises and extends into ER-PM contact sites. It traps and nucleates Orai1 (orange) into functional CRAC channels (step 2). For simplicity, Stim1 and Orai1 are illustrated as monomeric, but upon activation, Stim1 and Orai1 are proposed to oligomerise into dimers and hexamers, respectively. RHBDL2 (blue) cleaves the fourth transmembrane domain in Orai1, which anchors the primary carboxy-terminal Stim1-interaction site. **B**. Store-operated Ca^2+^ entry is monitored by cytosolic Fura-2 fluorescence, and compared between control HEK293 cells and those transiently transfected with GFP-RHBDL2 or Orai1 siRNA. Cells were stimulated with 2 mM thapsigargin in Ca^2+^ free buffer, followed by readmission of 1 mM external Ca^2+^, as indicated. Aggregate data from cells treated as in **B** are plotted, analysing the peak Ca^2+^ level in each condition (**C**) and rate of Ca^2+^ entry (**D**). Each bar in C and D represents between 34 and 68 cells. **E**. Ba^2+^ entry is compared between cells transfected with empty vector or GFP-RHBDL2, after treatment with 2 mM thapsigargin in Ba^2+^/Ca^2+^ free buffer. **F**. The rate of Ba^2+^ entry is plotted, each bar represents between 12 and 19 cells. For two-tailed t-tests *** = p<0.001, in comparisons with empty vector controls. In all bar charts, error bars indicate standard error of the mean. **G**. PFA-fixed HEK293 cells transfected with NFAT1(1-460)-GFP and indicated RHBDL2/4 constructs were treated with DMSO or 1 mM thapsigargin for 45 minutes. Single confocal sections of EGFP fluorescence are depicted with inverted grayscale lookup tables. Under each image, the number of cells displaying nuclear enriched NFAT-GFP is indicated. Scale bar = 10 μm.

RHBDL2 expression inhibits CRAC channel signalling, but is cleavage of Orai1 a physiologically meaningful event? To address this, we tested whether loss of RHBDL2 function had a physiological effect on Ca^2+^ signalling and the outputs of store-operated Ca^2+^ entry. We first used HEK293 mutant cells, in which the region encoding the catalytic histidine in RHBDL2 was deleted by CRISPR-Cas9 editing (**Figure S2A, B**). These cells displayed reduced endogenous store-operated Ca^2+^ entry across a range of physiological extracellular Ca^2+^ concentrations (**Figure 3A-3E**). We also assayed the effect of RHBDL2 depletion in HaCaT keratinocytes (**Figure 3F**). Using two different siRNAs, there was a clear defect in store-operated Ca^2+^ entry when RHBDL2 was depleted (**Figure 3G-3I**), further demonstrating that RHBDL2 does indeed regulate endogenous CRAC channel activity.

**Figure 3:**
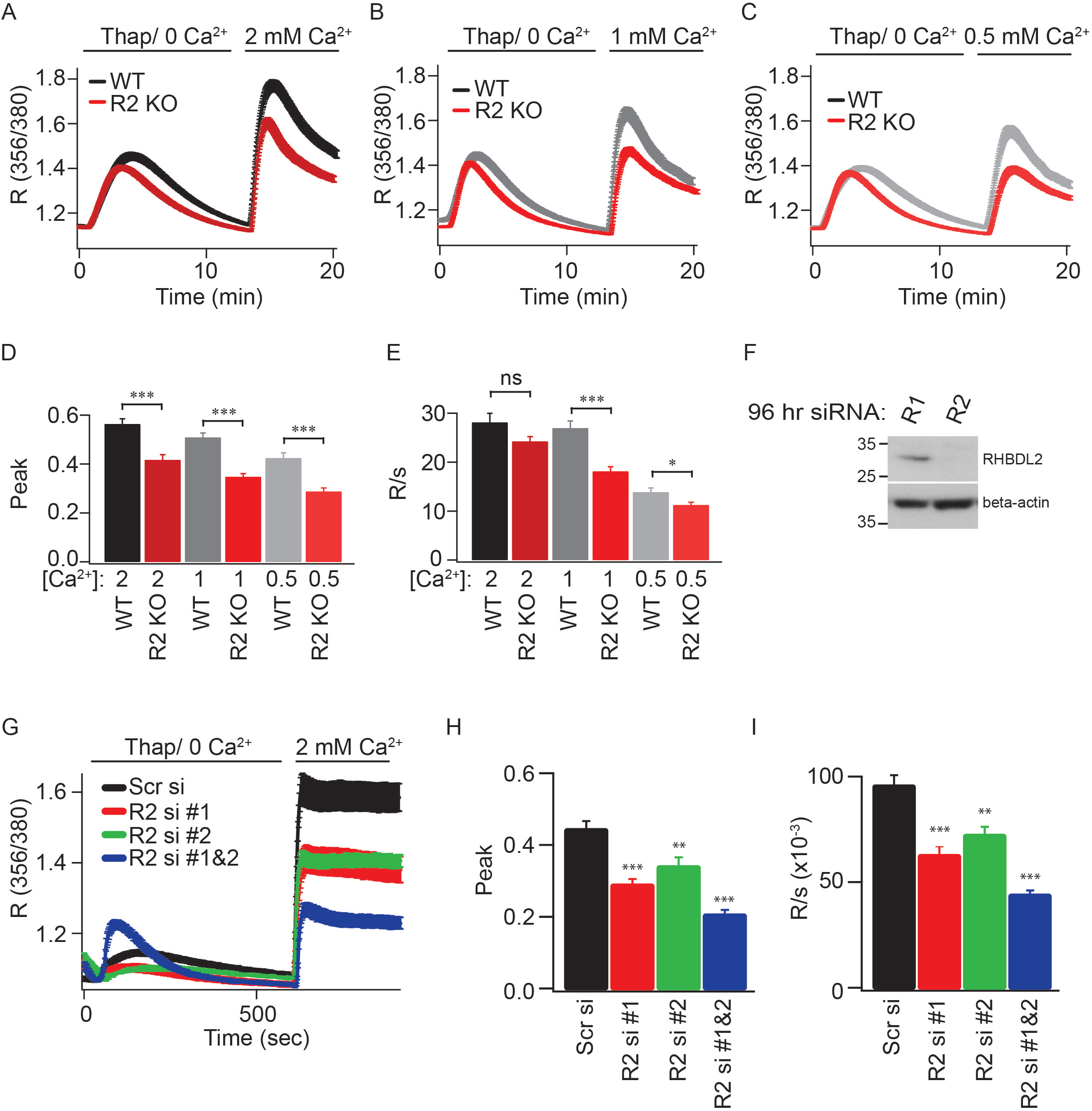
RHBDL2 is required for normal store-operated Ca^2+^ entry. **A-C.** Store-operated Ca^2+^ entry is monitored by cytosolic Fura-2 fluorescence, and compared between wild type and RHBDL2 mutant HEK293 cells (CRISPR/Cas9-based deletion of the essential catalytic histidine, termed R2 KO). Cells were stimulated with 2 mM thapsigargin in Ca^2+^ free buffer, followed by readmission of 2, 1 or 0.5 mM external Ca^2+^, as indicated. Aggregate data from cells treated as in **A-C** are plotted, analysing the peak Ca^2+^ level in each condition (**D**) and rate of Ca^2+^ entry (**E**). Each bar in **D** and **E** represents between 11 and 35 cells. **F**. Western blots of HaCaT keratinocytes treated with RHBDL1 or RHBDL2 siRNAs for 96 hours, probed with RHBDL2 and beta-actin antibodies. **G**. Store-operated Ca^2+^ entry is monitored via cytosolic Fura-2 fluorescence, and compared between scrambled siRNA and RHBDL2 siRNA-treated cells. Cells were stimulated with 2 mM thapsigargin in Ca^2+^ free buffer, followed by readmission of 2 mM external Ca^2+^. Aggregate data from cells treated as in **G** are plotted, analysing the peak Ca^2+^ level in each condition (**H**) and rate of Ca^2+^ entry (**I**). Each bar in H and I represents between 22 and 36 cells. For two-tailed t-tests * = p<0.05, ** = p<0.01 and *** = p<0.001, in comparisons with wild type or scrambled siRNA controls. In all bar charts, error bars indicate standard error of the mean.

### RHBDL2 is required for human T cell activation

CRAC channels participate in the primary activation of T cells by antigen presenting cells, and are thus centrally involved in T cell immunity (Feske, 2007). We therefore isolated primary CD4-positive T cells from two healthy human donors and asked whether RHBDL2 depletion (**Figure 4A and Figure S3A**) affected their activation by anti-CD3 crosslinking of the T cell receptor, a widely used method of mimicking T cell interactions with antigen presenting cells. Using surface CD69 expression as a readout (Ziegler et al., 1994), we found T cell activation was reproducibly defective across three experiments (**Figure 4B and Figure S3B;** *EC50 values of anti-CD3 for each shRNA are indicated in the dashed box*). We also directly measured CRAC channel activity in these RHBDL2-depleted T cells and found severely reduced store-operated Ca^2+^ entry **(Figure 4C-4E**). Combined, our results not only demonstrate that the role of RHBDL2 in controlling CRAC channel activity is essential for normal store-operated Ca^2+^ entry, but also that this has a profound effect on human T cell activation.

**Figure 4:**
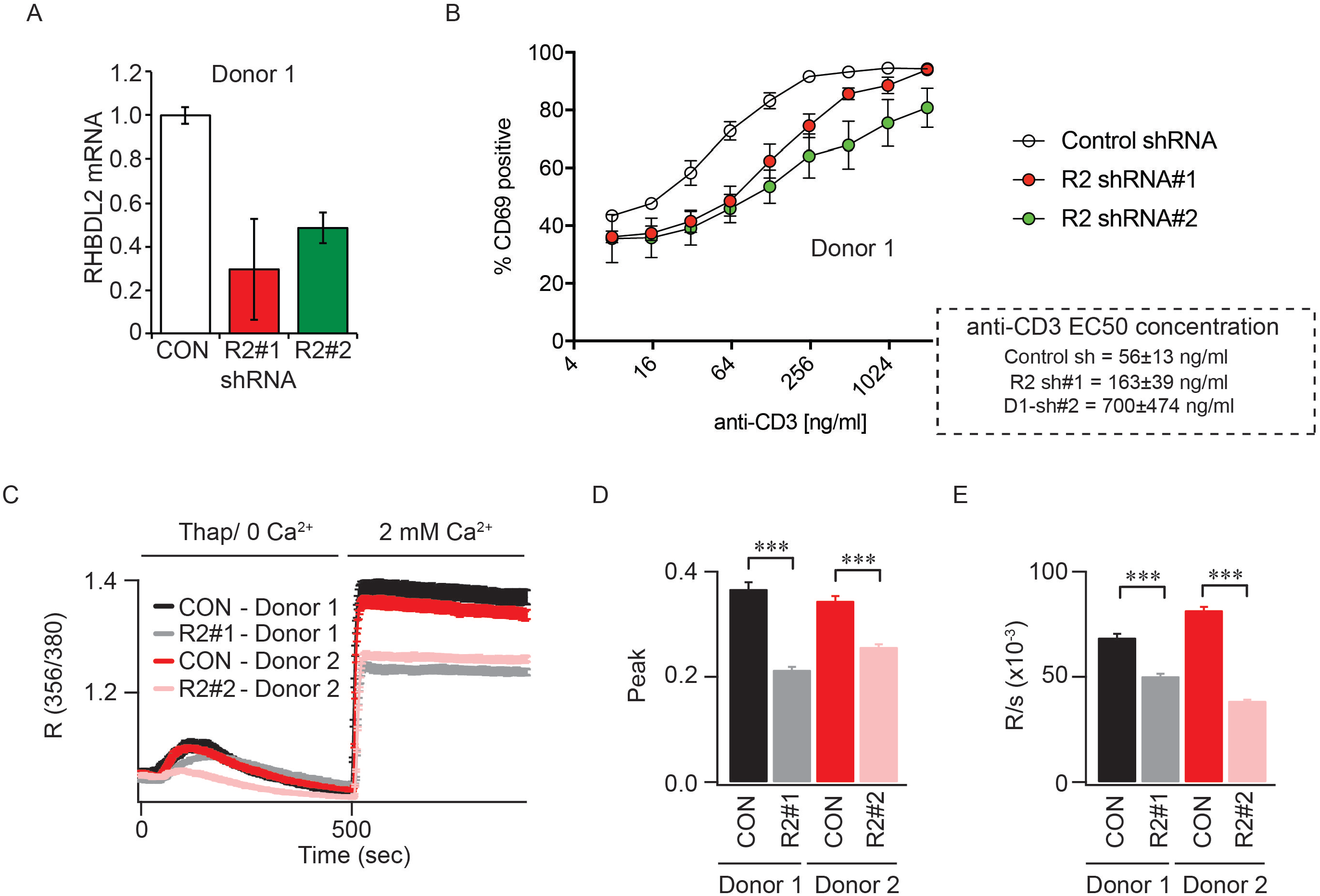
RHBDL2 is required for human T cell activation. **A**. TaqMan assays for RHBDL2 mRNA levels in T cells transduced with virus encoding control or RHBDL2 shRNAs. Error bars represent RQ standard error. **B.** T cell activation was measured by quantification of surface CD69 expression by FACS. CD69 expression is compared between control and RHBDL2 shRNA transduced primary CD4-positive T cells after stimulation with varying doses of platebound CD3. Each trace represents three biological replicates. In the dashed box, the calculated EC50 of anti-CD3 for each shRNA condition is indicated. Error indicates standard error of the mean. **C**. Store-operated Ca^2+^ entry is monitored by cytosolic Fura-2 fluorescence, and compared between control and RHBDL2 shRNA transduced T cells. T cells were stimulated with 2 mM thapsigargin in Ca^2+^ free buffer, followed by readmission of 2 mM external Ca^2+^. Aggregate data from T cells treated as in **C** are plotted, analysing the peak Ca^2+^ level in each condition (**D**) and rate of Ca^2+^ entry (**E**). Each bar in D and E represents between 34 and 45 cells. For two-tailed t-tests *** = p<0.001, in comparisons with control shRNA transduced T cells. Error bars represent standard error of the mean.

### RHBDL2 controls signalling by optimising stoichiometry between Orai1 and Stim1

Superficially, the loss of RHBDL2 might be expected to lead to higher Orai1 levels, more CRAC channels and therefore enhanced store-operated Ca^2+^ entry. We therefore sought to explain the counterintuitive result that loss of RHBDL2 led to decreased store-operated Ca^2+^ entry. We began by analysing the subcellular localisation of Orai1 in RHBDL2-depleted cells (**Figure 5A**). This experiment provided three important insights. First, Orai1 targeting to the PM was unaffected. Second, Orai1 levels at the PM appeared elevated in RHBDL2-depleted cells. This was further confirmed by cell surface biotinylation experiments, which showed a specific elevation of PM Orai1 levels upon depletion of RHBDL2 (**Figure S4**). Combined, this ruled out the possibility that the store-operated Ca^2+^ entry phenotype was due to a failure in trafficking of Orai1 to the PM. And third, we found that Orai1 did not accumulate in LAMP1-positive lysosomes, confirming that this increased pool of Orai1 at the PM upon knockdown of RHBDL2 was not a secondary consequence of defective lysosomal function. Consistent with its role in cleaving and targeting Orai1 for degradation, depletion of RHBDL2 – but not other rhomboids RHBDL1, 3 or 4 – led to an increase in endogenous full length Orai1 and its shorter isoform, Orai1β (**Figure 5B, 5C**). Plasmid-borne Orai1 also accumulated specifically in RHBDL2 depleted cells, clearly demonstrating that transcriptional changes are not responsible for increased Orai1 protein (**Figure 5D**). Overall, these data indicate that RHBDL2 loss caused elevated levels of PM Orai1, but further prompted the question of how this led to decreased store-operated Ca^2+^ entry.

**Figure 5:**
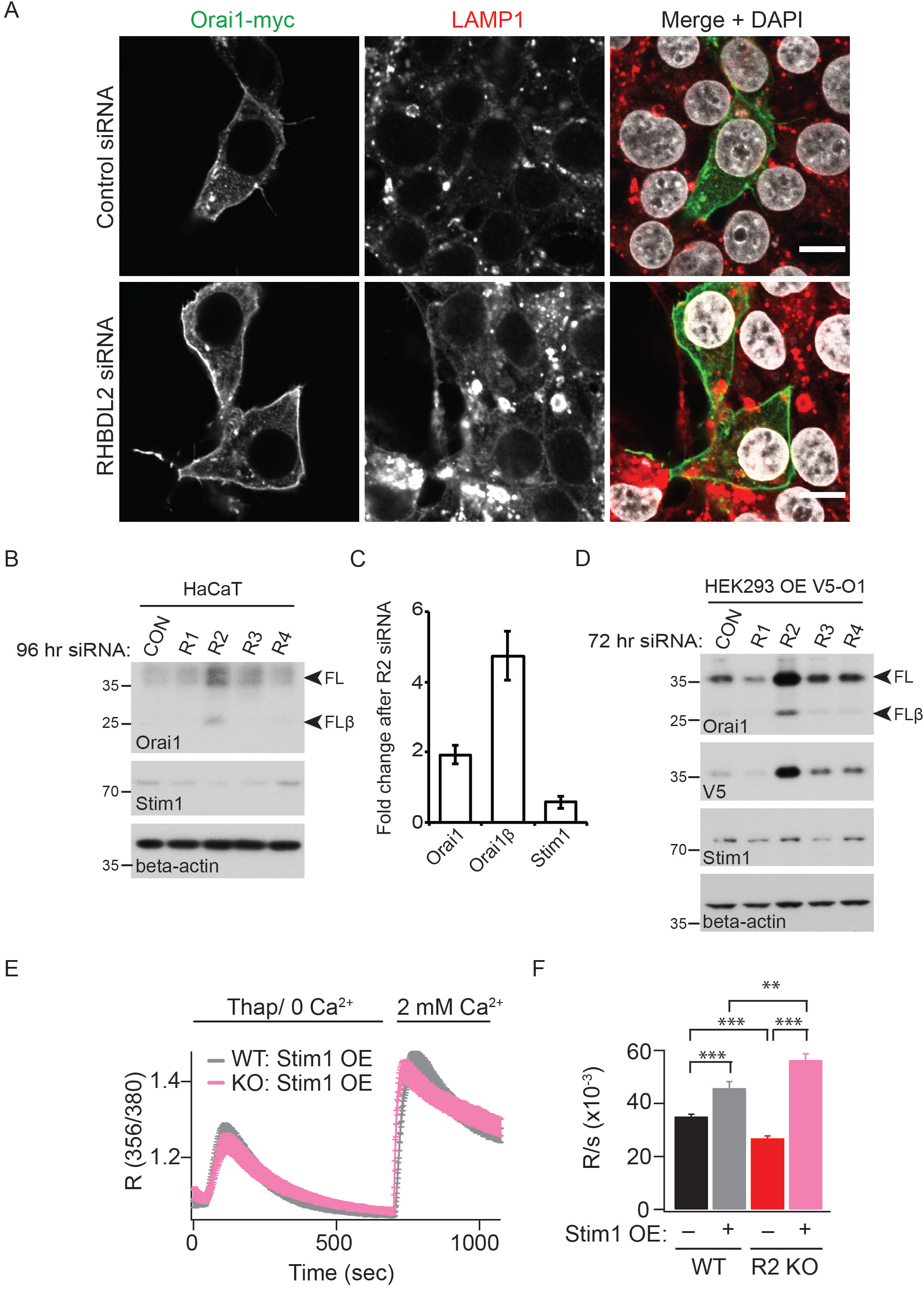
RHBDL2 controls signalling by optimising stoichiometry between Orai1 and Stim1. **A**. Immunofluorescent labelling of Orai1-myc and LAMP1 (to mark lysosomes) in HEK293 cells transfected with control or RHBDL2 siRNA for 72 hours, and transfected with Orai1-myc 24 hours prior to fixation. Individual confocal sections through the nucleus are depicted. **B**. Western blots of HaCaT lysates after cells were treated with control or RHBDL1-4 siRNAs for 96 hours, labelled for endogenous Orai1, Stim1 and beta-actin. Full length Orai1 (FL) and Orai1β (FLβ) are indicated by arrowheads. Orai1β arises from alternative start sites methionine-64 or -71 (Fukushima et al., 2012). **C**. Quantification of the fold change in Orai1 and Stim1 protein abundance, from three independent experiments performed as in B. Error bars represent the standard error of the mean. **D**. Western blots of HEK293T lysates after cells were treated with control or RHBDL1-4 siRNAs for 72 hours, expressing V5-Orai1 for the final 24 hours. Full length Orai1 (FL) and Orai1β (FLβ) are indicated by arrowheads. **E**. Store-operated Ca^2+^ entry is compared between wild type (WT) and RHBDL2 mutant HEK293 cells (CRISPR/Cas9-based deletion of the essential catalytic histidine, KO) over-expressing Stim1-YFP. Cells were stimulated with 2 mM thapsigargin in Ca^2+^ free buffer, followed by readmission of 2 mM external Ca^2+^. Aggregate data from cells treated as in E are plotted, analysing the rate of Ca^2+^ entry (**F**). Each bar in F represents between 17 and 26 cells. For two-tailed t-tests ** = p<0.001, *** = p<0.001, in comparisons with wild type cells. Error bars represent standard error of the mean.

The answer to this question was provided by the fact that the correct stoichiometry between Orai1 and Stim1 is essential for the store-operated Ca^2+^ entry pathway (**Figure 2A**): Stim1 levels are rate limiting for CRAC channel activation, and excess Orai1 has a dominant-negative effect on store-operated Ca^2+^ entry (Mercer et al., 2006; Peinelt et al., 2006; Soboloff et al., 2006; Scrimgeour et al., 2009; Hoover and Lewis, 2011;; Yeh et al., 2019). We found that RHBDL2 depletion caused PM Orai1 levels to increase, with unchanged or decreased Stim1 levels (**Figure 5B-5D**). We therefore predicted that if compromised stoichiometry was the cause of the observed store operated Ca^2+^ entry defects, overexpression of Stim1 in RHBDL2 KO cells should rescue the phenotype by allowing production of functional CRAC channels and elevated store-operated Ca^2+^ entry. Accordingly, expression of Stim1 not only rescued the defect in store-operated Ca^2+^ entry, but significantly enhanced its rate compared to wild-type cells expressing the same construct (**Figure 5E, 5F**). This result indicates that the defects in store-operated Ca^2+^ entry caused by RHBDL2 loss are caused by an imbalance in Orai1:Stim1 stoichiometry. An important implication of this result is worth emphasising: the Orai1 that accumulates in the absence of RHBDL2 is functionally competent. This rules out another possible role for RHBDL2, that it might act in a misfolded protein quality control mechanism to degrade defective Orai1, thereby protecting the integrity of CRAC channels. Overall, these data show that RHBDL2 cleavage and subsequent degradation of Orai1 acts to maintain an optimal stoichiometry between Orai1 and Stim1.

### RHBDL2 prevents inappropriate CRAC channel activation in resting cells

We next questioned the biological context of Orai1 cleavage by RHBDL2. We noted that in unstimulated cells, PM Orai1 protein levels increased upon depletion of RHBDL2 (**Figure 5A and Figure S4**), indicating that cleavage was not dependent on store-operated Ca^2+^ entry. We therefore hypothesised that the role of RHBDL2 cleavage of Orai1 is to prevent CRAC channel activity in the absence of stimulation, i.e. that RHBDL2 maintains the correct baseline threshold level of CRAC channel signalling. NFAT nuclear translocation is highly sensitive to low-level local CRAC channel activation (Kar et al., 2011; Kar and Parekh, 2013). As NFAT translocation is prevented by RHBDL2 expression (**Figure 2G**), we reasoned that if RHBDL2 loss led to a basal elevation in CRAC channel activity in unstimulated cells, NFAT targets would be upregulated. Strikingly, we found that RHBDL2 depletion in HaCaT cells led to a 19-fold upregulation of the expression of the inflammatory cytokine TNF, one of the major NFAT responsive genes (Rao et al., 1997) (**Figure 6A;** *72 hours siRNA*). Notably, cytokines not dependent on NFAT, such as IL-6, were not affected. We confirmed that the elevated TNF expression was indeed due to upregulated NFAT, as it was inhibited by treatment with cyclosporin A, a widely used inhibitor of NFAT signalling (Rao et al., 1997) (**Figure 6B**). Together, these data demonstrate that RHBDL2 is needed to prevent inappropriate NFAT signalling, which is a major downstream effector of CRAC channels.

**Figure 6:**
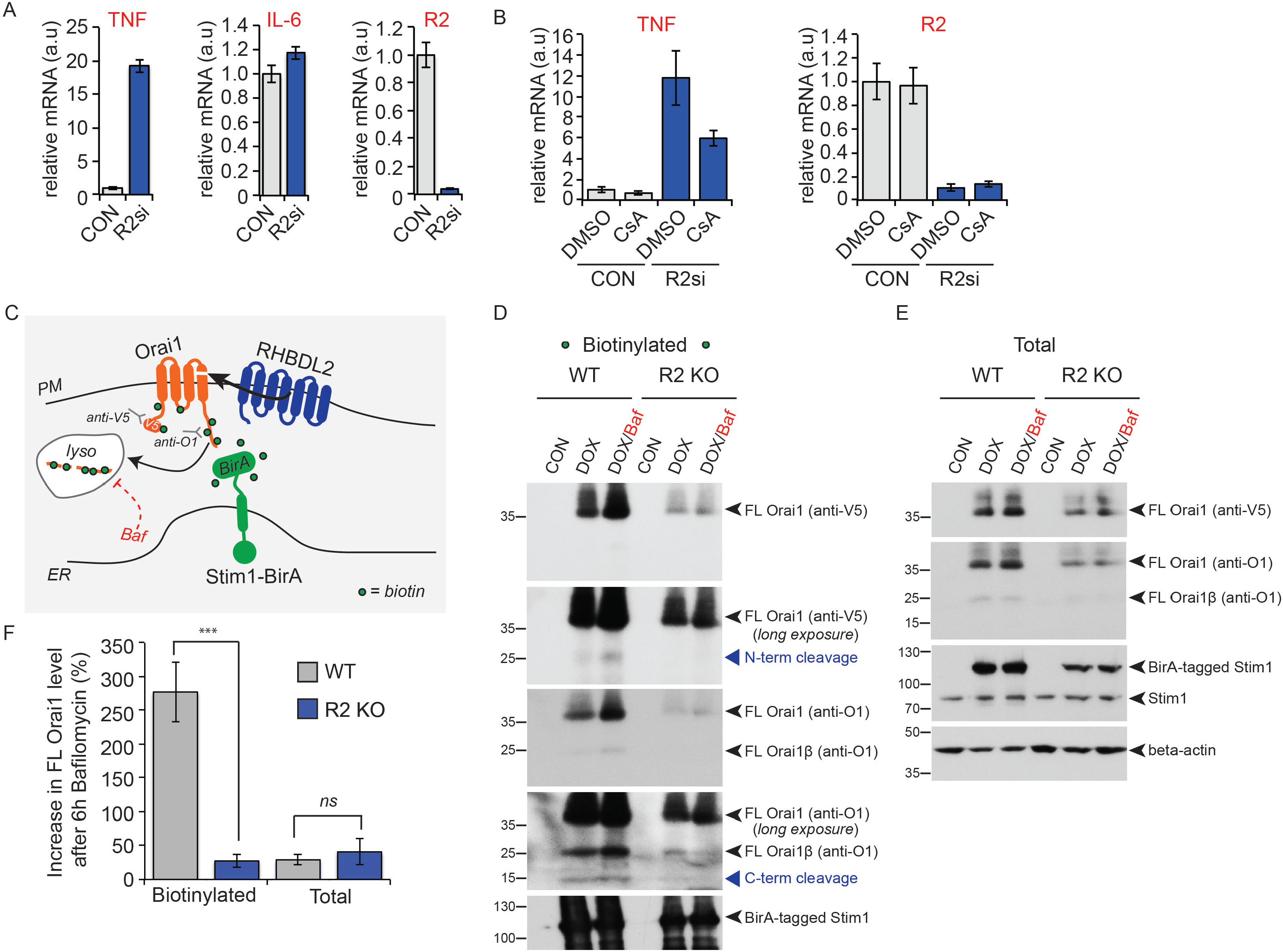
RHBDL2 prevents inappropriate CRAC channel activation in resting cells. **A-B**. TaqMan assays for TNF alpha, IL-6 and RHBDL2 (R2) mRNA levels in HaCaT cells treated with control or RHBDL2 siRNAs for 72 hours (in **A**) or 48 hours (in **B**). Cyclosporin A (1 μm) was added for the final 24 hours in B. Error bars represent RQ standard error. Each chart in A and B represents one of at least four biological replicates. **C**. A scheme of the Stim1-BirA experiment in **D-F**, illustrating the biotinylation of V5-Orai1 (orange) by Stim1-BirA* (green) at PM-ER contact sites, and the downstream consequence of RHBDL2 (blue) activity. The epitopes in Orai1 that are recognised by antibodies used in D-F are shown, as well as the inhibition of downstream lysosomal degradation by bafilomycin A1 (Baf). Biotin is indicated by green dots. For simplicity, Stim1 and Orai1 are illustrated as monomeric. **D-E**. Western blots of neutravidin agarose-based biotin captured lysates from wild type (WT) or RHBDL2 knockout (R2 KO) HaCaT cells. The expression of V5-Orai1 and Stim1-BirA* was induced with doxycycline (DOX, 250 μg/ml final) for 96 hours in the presence of 50 μm biotin. Six hours prior to lysis, bafilomycin A1 (BAF, 100 nm final) was added to block lysosomal degradation. Blots were probed for the N-terminal epitopes or C-terminal epitopes recognised by V5 and O1 antibodies, respectively. Stim1 and Stim1-BirA* were probed for using an anti-Stim1 antibody. Different full length forms of Orai1 (Orai1β arises from alternative start sites methionine-64 or -71 (Fukushima et al., 2012)) and their cleavage products are indicated by the blue arrowheads. **F.** Quantification of the increase in full length Orai1 upon bafilomycin treatment from three replicates of the experiment performed in **D-E.** Error bars indicate standard error of the mean. For two-tailed t-tests ns = not significant, *** = p<0.001, for indicated comparisons.

We next examined whether excess signalling was caused by stimulus-independent, stochastic Stim1 activation of CRAC channels. This was prompted by two observations. First, PM-ER contact sites stably exist regardless of Stim1 activation (Wu et al., 2006; Orci et al., 2009), and Stim1 targeting to PM-ER contact sites is Orai1-independent (Liou et al., 2007; Park et al., 2009). Second, Stim2, which is prelocalised at PM-ER contact sites, can promote Stim1 translocation in conditions of incomplete depletion of ER Ca^2+^ stores (Burdakov and Verkhratsky, 2006; Brandman et al., 2007; Subedi et al., 2018). We therefore hypothesised that the unwanted CRAC channel activity against which RHBDL2 protects cells may be stochastic, triggered by random Stim1/Orai1 interaction in the absence of stimulation, rather than being actively triggered by complete depletion of ER Ca^2+^. To test this idea, we fused the BirA* biotin ligase to the cytoplasmic domain of Stim1, the basis of an assay to identify Orai1 molecules that have previously encountered Stim1. We then asked whether this subset were preferentially cleaved by RHBDL2 (**Figure 6C**). At rest, in unstimulated wild type cells, a small proportion of Orai1 (0.38 ± 0.13% of the total pool after 72 hours of expression) does indeed encounter Stim1 (**Figure 6D;** *DOX*). Importantly, this pool of Orai1 was cleaved in wild-type cells in a RHBDL2-dependent manner (**Figure 6D**- *N- and C-terminal cleavage products in WT vs KO;* **Figure S2C-D**). Blocking lysosomal degradation increased these cleavage products. These cleavage products were never detected in total lysates, indicating the specificity of RHBDL2 for the small pool of Orai1 that had inappropriately encountered Stim1 (**Figure 6E**). The central message of this experiment is that Orai1 was cleaved by endogenous RHBDL2, and subsequently degraded in lysosomes, only after stimulus-independent engagement with Stim1.

Secondarily, we noted that full length Orai1 (i.e. uncleaved by RHBDL2) that had previously encountered Stim1-BirA* was also stabilised by bafilomycin treatment (**Figure 6D**), indicating it too was degraded in lysosomes. Intriguingly, this did not occur in cells lacking RHBDL2 (**Figure 6D, 6F**-*biotinylated O1 levels in WT versus R2 KO)*. One possible interpretation of this phenomenon is that cleavage of one or two copies of Orai1 is sufficient to destabilise larger homomeric Orai1 complexes, leading to degradation of both full-length and cleaved protein.

Overall, these results define a central conclusion of our work: that RHBDL2 promotes the cleavage and subsequent lysosomal degradation of only those Orai1 molecules that have previously encountered Stim1. In unstimulated cells, this population of molecules is small but, as shown in Figures 4B and 6B, they nevertheless trigger a dangerous level of unwanted Ca^2+^ signalling if allowed to accumulate. Without RHBDL2 acting as a brake on this stochastic CRAC channel activity, T cell activation and inflammatory cytokine expression are both severely defective.

### RHBDL2 recognition of Orai1 is conformationally determined

The proposed model of RHBDL2 patrolling the PM to destroy inappropriately active CRAC channels suggests that the protease may preferentially recognise active forms of Orai1, engaged by Stim1. Such conformational selectivity of rhomboid proteases has not previously been reported. To address the idea, we capitalised on recent structure-function data that provides a detailed understanding of the contributions of specific Orai1 TMD amino acids to overall CRAC channel architecture and activity (Yamashita et al., 2017; Hou et al., 2018). Mutation of histidine-134 in Orai1 to threonine, valine or serine activates Orai1 (Yeung et al., 2018), as do other mutations such as F99Y, V102A and P245L (Nesin et al., 2014; Palty et al., 2015). Conversely, other Orai1 TMD mutations have a profound inactivating effect (G98C (Yamashita et al., 2017), R91W (Feske et al., 2006)) and H134W (Yeung et al., 2018)). We compared binding of RHBDL2-SA (the serine-to-alanine catalytic mutant, which binds stably to substrates) to these different Orai1 activity mutants. There was a clear correlation between RHBDL2-SA binding and Orai1 activity: RHBDL2 bound strongly to Orai1 H134S, the mutant that is closest in its properties to a Stim1-gated CRAC channel (Yeung et al., 2018). In contrast, inactive mutants of Orai1, such as H134W, showed very weak binding to RHBDL2 (**Figure 7A, 7B**). This demonstrated that RHBDL2 exhibits selectivity for active forms of Orai1.

**Figure 7:**
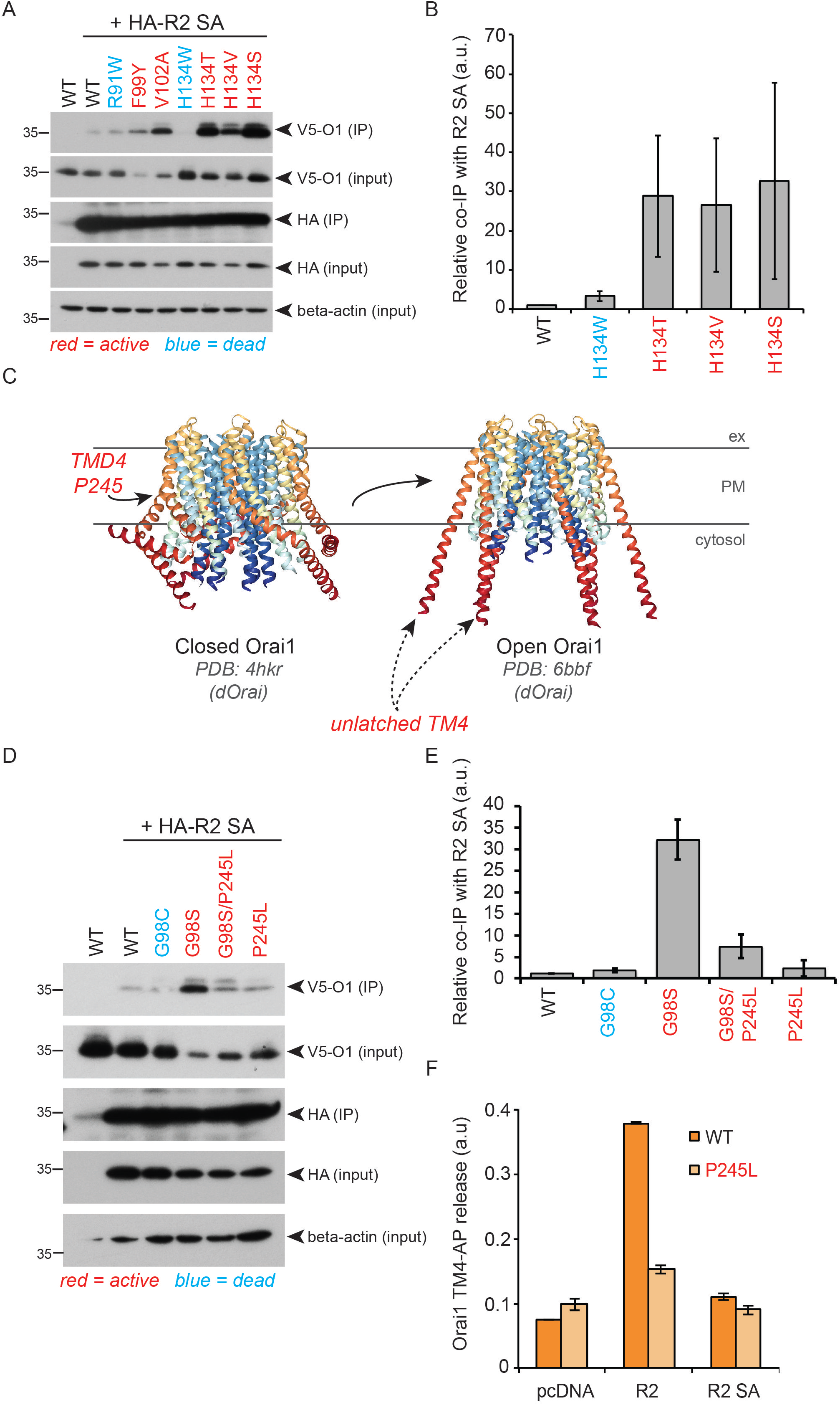
RHBDL2 recognition of Orai1 is conformationally determined. **A.** HA immunoprecipitates (IP) and inputs from HEK293 cells transiently expressing 3xHA-RHBDL2 SA and wild type or mutant V5-Orai for 24 hours were immunoblotted for V5, HA and beta-actin. **B**. Quantification of three biological replicates of the experiment performed in **A**. **C**. Structures of Drosophila Orai WT (left, PDB: 4hkr) or Orai H134A (right, PDB: 6bbf), highlighting the accessibility of the fourth transmembrane domain within the membrane, and the large change in conformation around P245 that is associated with CRAC channel activity. **D**. HA immunoprecipitates (IP) and inputs from HEK293 cells transiently expressing 3xHA-RHBDL2 SA and wild type or mutant V5-Orai for 24 hours were immunoblotted for V5, HA and beta-actin. **E**. Quantification of three biological replicates of the experiment performed in **D. F.** HEK293 cells were transfected with pcDNA, 3xHA-RHBDL2 or RHBDL2 SA, and AP-TMD4 or AP-TMD P245L from Orai1 for 48 hours. Released AP was collected over the final 16 hours. Values represent the level of released alkaline phosphatase/total alkaline phosphatase. Error bars represent standard error of the mean.

We next questioned whether RHBDL2 recognises the active form of Orai1 through recognition of activity-dependent conformational changes. The structure of the active *Drosophila* Orai1 H134A mutant also mimics the conformation of Orai1 in complex with Stim1, showing the major displacement of TMD4, which anchors the Stim1 interacting cytoplasmic domain (Hou et al., 2018) (**Figure 7C**). This activating displacement pivots on a flexible hinge generated by a proline residue in TMD4 (proline-245 in human Orai1) (Hou et al., 2012; Hou et al., 2018). Mutation of this hinge proline is also the cause of the rare human Stormorken syndrome, in which the CRAC channel has excess activity (Nesin et al., 2014). Since helical instability, often conferred by prolines, is a major determinant of rhomboid substrates (Urban and Freeman, 2003; Moin and Urban, 2012), we examined the role of proline-245 in RHBDL2 recognition of Orai1. Unlike all other active Orai1 mutants we tested, P245L did not show enhanced binding to RHBDL2-SA, indicating that, even when the molecule is in a locked-open state, proline-245 is necessary for RHBDL2 recognition (**Figure 7D**). Moreover, when combined with the activating G98S mutation (Endo et al., 2015), which itself strongly promotes RHBDL2 binding, the P245L mutation prevented recognition (**Figure 7D, 7E**). Finally, RHBDL2 proteolysis of Orai1 TMD4 was inhibited by the P245L mutation (**Figure 7F**), confirming that RHBDL2 recognition and cleavage of Orai1 requires proline-245. This demonstrates that loss of helical instability in TMD4, which activates Orai1 and causes Stormorken syndrome, also blocks recognition by rhomboid and thus prevents its ability to perform conformational surveillance.

## Discussion

The results we report here uncover a new role for rhomboid proteases in regulated degradation of membrane proteins. RHBDL2 patrols the PM to seek and destroy inappropriately activated CRAC channels in unstimulated cells, in order to prevent inappropriate signalling such as NFAT-dependent expression of the proinflammatory cytokine, TNF. This maintains a low baseline level of Ca^2+^ influx, which is essential to ensure fully regulated switch-like store-operated Ca^2+^entry and T cell activation. It is notable that RHBDL2 has been shown to diffuse in the plane of the membrane at exceptionally high speeds, faster than any known polytopic membrane protein (Kreutzberger et al., 2019). This property, which appears to be mediated by mismatch between the thickness of the lipid bilayer and the shorter than expected hydrophobic domain of the rhomboid fold, makes RHBDL2 particularly well suited to this seek-and-destroy function. We discovered Orai1 using a combined bioinformatic and cell-based screen for new substrates of the PM rhomboid RHBDL2. This approach identified specifically the fourth TMD of Orai1 (as well as Orai2 and Orai3). Combined with the recent discovery of a polytopic substrate for the bacterial rhomboid protease YqgP (Began et al., 2020), the universe of possible rhomboid substrates has therefore been expanded beyond single pass TMD proteins, to now encompass the very large class of polytopic membrane proteins, which includes channels, GPCRs and many other pharmacologically significant targets.

Prolines are often determinants of rhomboid substrates, because of their property of destabilising or introducing a kink into a transmembrane helix (Urban and Freeman, 2003). This partial disruption is needed to allow the active site of the enzyme access to the cleavable peptide bond, which is otherwise shielded by hydrogen bonding inherent to the alpha helix. Our data elaborate on this basic principle of rhomboid substrate recognition by identifying the first case in which a rhomboid, or indeed any intramembrane protease, shows conformational specificity in substrate recognition. RHBDL2 monitors the conformational dynamics of Orai1. Proline-245 of Orai1 contributes the flexibility to the fourth TMD that is essential for transducing the binding of Stim1 into the allosteric changes that open the CRAC channel. This same proline-245 hinge mechanism determines recognition by RHBDL2. The ability to distinguish active from inactive Orai1 underlies the ability of RHBDL2 to seek and cleave only activated CRAC channels, and is therefore central to the mechanism of maintaining low basal signalling in unstimulated cells. Interestingly, transmembrane helix instability is a common characteristic of most intramembrane protease substrates (Ye et al., 2000; Lemberg and Martoglio, 2002; Urban and Freeman, 2003), which raises the possibility that other intramembrane proteases may perform a similar function.

Our model that RHBDL2 acts to prevent inappropriate CRAC channel activity, begs the question of how stimulated activity occurs when store operated Ca^2+^ entry appropriately triggers signalling. What prevents RHBDL2 from blocking signalling in a scenario when signalling is needed? We propose three possible answers to explain this. First, store operated Ca^2+^ entry leads to molecular crowding of CRAC channels in the membrane, with an estimated ~40 nm distance between channels (Ji et al., 2008). Such a high density of CRAC channels may physically restrict RHBDL2 access to substrate TMDs within the plane of the membrane. Second, in unstimulated cells only a small proportion of Orai1 encounters Stim1, allowing the low level of RHBDL2 expressed in most cells to be sufficient to prevent unstimulated signalling (our observations, and https://gtexportal.org/home/). In contrast, when store-operated Ca^2+^ entry is triggered, the majority of Orai1 is engaged, and this may simply overwhelm RHBDL2 surveillance. A third more speculative possibility is that when CRAC channels are activated, RHBDL2 function is inhibited by an increase in cytoplasmic Ca^2+^. Significantly, there is precedent for rhomboid proteases being Ca^2+^-sensitive (Baker and Urban, 2015), so this could in principle provide a neat regulatory mechanism to prevent Orai1 cleavage when store operated Ca^2+^ entry is triggered.

CRAC channel activity is a major mechanism for regulating cytoplasmic Ca^2+^ levels in non-excitable cells and therefore plays an important role in a wide range of biological contexts, most notably during the activation of T cells when they engage with antigen presenting cells (Feske, 2007). Although store-operated Ca^2+^ entry, dependent on phospholipase C activity, is a tightly regulated process, even highly evolved biological control processes are not perfect. The events downstream of CRAC channels are biologically potent and, if unchecked, they cause pathophysiological dysregulation and channelopathies (Feske, 2010). Accordingly, our data show that the loss of RHBDL2 from cells leads to significantly elevated levels of Orai1, and consequent dysregulated T cell activation and inflammatory cytokine expression. Significantly, the major determinant of RHBDL2 recognition of Orai1, proline-245, is the causative mutation of the rare inherited Stormorken Syndrome, which is characterised by excess CRAC channel activity (Nesin et al., 2014). The aetiology of this disease, and perhaps others caused by excess CRAC channel activity, is therefore likely to be related to failure of RHBDL2 surveillance of Orai1.

In conclusion, the identification of Orai1 as a substrate of RHBDL2 highlights two novel themes. It substantially advances our knowledge of rhomboid proteases by expanding the universe of potential substrates, and by demonstrating the first example of intramembrane proteases showing conformation-specific substrate recognition, which has significant implications for their regulatory roles. Our work also develops a theme of regulated protein degradation being used to sharpen cellular signalling, by ensuring low levels of activity in unstimulated cells. RHBDL2 patrols the PM, seeking Orai1 molecules in an inappropriately active conformation in resting cells, and triggering their degradation. Rhomboid proteases are ancient, existing in all kingdoms of life. It is tempting to speculate that the primordial function of rhomboids may have been to inactivate and degrade membrane proteins with non-canonical TMDs, perhaps as a quality control function. In this scenario, it would only have been later, after the appearance of metazoans, when rhomboids would become responsible for their now well established roles in triggering the release of proteins that signal between cells.

## Acknowledgements

We thank members of the Freeman lab for their support and advice during the study, particularly Fangfang Lu and Nina Jajcanin-Jozic for their feedback on the manuscript. We thank Pedro Carvalho (Dunn School, Oxford), Tim Levine (Institute of Ophthalmology, UCL) and Francesca Robertson (Biochemistry Department, Oxford) for critical reading of the manuscript. We also thank Tim Levine for his guidance with the use of HHpred. We would like to thank Michael van der Weijer for the generous gift of pLVX plasmids subcloned with zeocin and blasticidin resistance genes. We gratefully acknowledge the support of Alan Wainman in the light microscopy facility and Michal Maj in the flow cytometry facility at the Dunn School. This paper was supported by the following grants: CIU Trust grant to JB. European Regional Development Fund (ERDF/ESF; project no. CZ.02.1.01/0.0/0.0/16_019/0000729) to KS. MRC grant (LO1047CX) to ABP. Marie Skłodowska-Curie fellowship (Horizon 2020 Framework Programme 659166) to AGG. BBSRC Research grant (BB/RO16771/1) to AGG and MF. Wellcome Trust Senior Investigator Award (101035/Z/13/Z) to MF.

## Author Contributions and Declarations of Interest

YY was responsible for design and implementation of all Fura-2/calcium experiments, YY and ABP analysed these data. LZ contributed to the RHBDL2/Orai1 binding assays. JB conducted all T cell activation assays, and MHB provided guidance in T cell work its interpretation. NJ and KS made and provided HaCaT RHBDL2 KO keratinocytes. AGG led the project and performed all other experiments. AGG and MF conceived the project and wrote the manuscript.

The authors have no conflict of interest.

## Experimental procedures

### Reagents

Bafilomycin A1 (catalogue number 19-148), biotin (catalogue number B4639), blasticidin (catalogue number 15205) and thapsigargin (catalogue number T9033) was purchased from Sigma Aldrich. Doxycycline was purchased from MP Biomedicals (catalogue number SKU 0219504405). PNGase F was purchased from New England Biolabs (catalogue number P0704L). Puromycin was purchased from Gibco (catalogue number A11138-03). Zeocin was purchased from Invitrogen (catalogue number 2058442). Phosphatase substrate kits containing PNPP tablets and buffer (catalogue number 37620) were purchased from Thermo Scientific.

### Antibodies

The following antibodies were used for western blotting (WB) and immunofluorescence (IF): mouse anti-beta-actin (Santa Cruz, catalogue number sc-47778; WB 1:2000), mouse anti-HA (ENZO, catalogue number ENZ-ABS120-0200; WB 1:1000), mouse anti-transferrin receptor (Invitrogen, catalogue number 13–6800; WB 1:1000), rabbit anti-Stim1 (Cell Signalling Technology, catalogue number 5668S (D88E10); WB 1:2000), rabbit anti-Orai1 (Sigma Aldrich, catalogue number O8264; WB 1:2500), goat anti-myc tag (Abcam, catalogue number ab9132; IF 1:2000), rabbit anti-RHBDL2 (Proteintech, catalogue number 12467-1-AP; WB 1:250 – only detected RHBDL2 in HaCaT lysates), rabbit anti-V5 tag (Cell Signalling Technology, catalogue number 13202S; WB and IF 1:2000). Corresponding species-specific HRP or fluorescently coupled secondary antibodies were used from Santa Cruz and Cell Signaling (WB) or Invitrogen (IF).

### Molecular biology

For generation of the AP reporter construct, we PCR amplified sequence encoding the signal sequence of HB-EGF and alkaline phosphatase, as described originally in (Sahin et al., 2004), with a pair of restriction enzyme sites (SalI and NotI) that were placed 3’ of the sequence encoding alkaline phosphatase. This was subsequently cloned between the EcoRI and SalI sites in the multiple cloning site of pcDNA3.1. This created a construct (pcDNA3.1_TMDscreen) that expresses a protein that constitutively enters the endoplasmic reticulum. Sequences encoding candidate TMDs, plus 3 cytoplasmic amino acids and 7 extracellular amino acids, were then ordered as paired oligonucleotides with 20 base pairs of overlap, and with overhangs that complemented the SalI and NotI sites in pcDNA3.1_TMDscreen. Paired oligonucleotides were extended on one another with 3 rounds of 98°C (15 sec) → 55°C (20 sec) → 72°C (30sec), followed by 72°C (7 min) in a thermal cycler. Double stranded oligos were then column purified and cloned into SalI-NotI digested pcDNA3.1_TMDscreen by InFusion cloning, test digested and positive colonies were confirmed by Sanger sequencing (Source Bioscience). pcDNA3.1 vectors encoding 3×HA-mRHBDL1-4 and GFP-mRHBDL2 have been previously described elsewhere (Lohi et al., 2004; Adrain et al., 2011). Inactive S->A mutants, and AP-TMD4 Orai1 point mutant were generated using site directed mutagenesis kits, according to the manufacturers instructions (Agilent). Stim1-YFP and GFP-NFAT1 (1-460) were previously used and described in Kar et al., 2011. Orai1-myc was previously described in Yeh et al., 2019. To generate stable HaCaT cell lines inducibly expressing V5-Orai1 and Stim1-BirA we used pLVX plasmids (Takara) that were subcloned to express zeocin and blasticidin resistance genes, a kind gift from Dr Michael van der Weijer (Dunn School, Oxford). To generate pcDNA3.1-V5-Orai1 and pLVX-V5-Orai1, we PCR amplified human Orai1 either flanked by the regions surrounding the BamHI/XhoI sites in pcDNA3.1 or the AgeI site in pLVX_Blasticidin. After digestion of pcDNA3.1 and pLVX_Blasticidin with corresponding restriction enzymes, we then inserted V5-Orai1 by InFusion cloning, according to manufacturers instructions (Takara). To generate pLVX-Stim1-BirA, we PCR amplified human Stim1 from Stim1-YFP, and BirA* (Roux et al., 2012), with 20 nucleotides of overlap with one another and the sequence flanking the AgeI site in pLVX_Zeocin. After digestion of pLVX_Zeocin with AgeI, we then inserted Stim1-BirA* by InFusion cloning, according to the manufacturers instructions (Takara).

### Cell culture

All cells used were maintained in regular high-glucose DMEM, supplemented with 10% FCS, 100μg/ml penicillin, and 100μg/ml streptomycin. All cells used in this study were subject to regular mycoplasma testing.

### CRISPR-Cas9 gene editing

For all editing, sequences of suitable guide RNAs were designed using publicly available prediction tools at www.broadinstitute.org/rnai/public/analysis-tools/sgrna-design (Doench et al., 2014) and at http://crispr.mit.edu/ (Hsu et al., 2013). A paired nickase Cas9 strategy was used to target the catalytic histidine in HEK cells to generate RHBDL2 KO HEKs. Guides targetting the following loci in human chromosome 1: TGAGCTGCAAAAGACACCTTGGG(−) and GGATTTGCTGGAATGTCCATTGG(+) were chosen. Guide sequences, without the PAM, were cloned into pSpCas9n(BB)-2A-Puro (px462), and sequence verified. Cells grown in 6 well dishes at 30-40% confluency were transfected with 500 ng CRISPR/Cas9. 24 h later cells were selected in 1μg/ml puromycin (Gibco) overnight. After 24 h recovery, 100 cells were seeded for colony growth in 10cm dishes. Colonies were picked used cloning discs and cells were amplified in 24 well dishes. Expanded colonies were then lysed at 65°C in 10 mM Tris-HCl (pH 8), 25 mM NaCl, 1mM EDTA, and 200 μg/ml proteinase K. Proteinase K was inactivated at 95°C for 2 min and samples were analysed via PCR and high resolution melt analysis, according to (Bassett et al., 2013), and deletions were confirmed by Sanger sequencing (Source Bioscience, Oxford). Clone “5j” was found to contain a deletion around the catalytic histidine, which would also produce a premature stop codon. To target the endogenous RHBDL2 gene in HaCaT cells using CRISPR/Cas9 we introduced a premature stop codon within exon 2 of the endogenous RHBDL2 gene by introducing indels targeting a site 115 bp after the initiator ATG codon of RHBDL2 (**Figure S2C**). Two of the four highest scoring guide RNA sequences (gRNA4: CCAAGAGTAAAAAGGTCCAC and gRNA1: ATGCTGCCCGAAAAGTCCCG) were cloned into pLenticrisprv2 (Sanjana et al., 2014) to yield targeting constructs pPR62 and pPR63, respectively. HaCaT cells were seeded at 5×10^5^ cells per 6 cm dish. The next day cells were transfected with 2 μg pPR62 or pPR63 using Fugene 6. 24 h later cells were selected using 2 μg/ml puromycin for 96 hrs. Then media was exchanged for complete DMEM and cells were allowed to recover to confluence, at which point they were trypsinized, suspended in 1 ml PBS + 2% FBS and single cell sorted into 96 well plates containing 20% FBS, 50% conditioned medium, 30% complete medium and 1×Gentamycin/AmphotericinB (1 μg/ml and 250 ng/ml, respectively; Thermo R01510). Single cells were allowed to proliferate and were expanded for analysis by indel screening and western blotting. To screen for indels, genomic DNA from each clone was isolated, amplified by PCR using primers flanking the Cas9 cleavage site, and products were analysed by TBE agarose electrophoresis and comparison to untargeted HaCaT cells. Positive clones containing indels and lacking the wild type allele were further characterised by DNA sequencing and western blotting. For sequencing, primers flanking the Cas9 cleavage site were used to amplify this region by PCR from the genomic DNA of candidate KO clones before cloning into a vector, transformation and propagation in *E.coli*. A minimum of 7 colonies per clone were sequenced to identify all possible genomic alterations at the Cas9 cleavage site in the hypotetraploid HaCaT cells. Ultimately, absence of endogenous RHBDL2 protein and enzymatic activity was verified by western blotting using a polyclonal antibody against human RHBDL2 and an RHBDL2 substrate (**Figure S2D**). To serve as controls, untargeted wild type HaCaT cells were single-cell sorted and expanded into clonal cultures. Wild type clone B10 and RHBDL2 deficient clone H9 (generated with gRNA4) were used in all experiments in Figure 6C-F.

### Lentivirus production and transduction

HEK293 cells grown to 30-40% confluence in 6-well dishes were transfected with Lipofectamine 2000 (Invitrogen) in 35mm plates with 0.5 μg of pLKO shRNA or pLVX expression plasmids (Takara), 0.35 μg pCMV-dR8.2 and 0.15 μg pCMV-VSVG. The pLKO shRNA plasmids (Control shRNA against RhoGDI: CA143; RHBDL2 #1: CA146; RHBDL2 #2: CA148; RHBDL2 #3: CA149) were previously validated by (Adrain et al., 2011). The following day, medium was changed and transfected cells were allowed to secrete virus for 48-72 hours in 2 ml complete medium. Culture supernatants were then centrifuged clarified by filtration with Sartorius Minisart syringe filters (0.45 μm pore size). For infection of HaCaTs or primary CD4-positive T cells, cells were split the day before, and viral supernatants were diluted 1- or 2-fold in fresh medium for transduction. Transduction was carried out in the presence of 10 μg/ml polybrene and a medium change was made 24 hours later. For selection, cells were treated with 10 μg/ml puromycin, 100 μg/ml zeocin or 5 μg/ml blasticidin, until all cells were killed in control transductions. In the case of the Stim1-BirA* HaCaTs, WT and KO cells were first transduced with pLVX-V5-Orai1-myc (and selected with blasticidin), followed by a second transduction with pLVX-Stim1-BirA* (and selected with zeocin).

### siRNA

Orai1 siRNA was purchased from Horizon, ON-TARGET SMARTPool (Catalog ID: L-014998). A final concentration of 50 nM was used to knockdown Orai1. The following siRNAs against human rhomboid proteases were purchased from Invitrogen: RHBDL1 #1: HSS113329; RHBDL1 #2: HSS113330; RHBDL2 #1: HSS123556; RHBDL2 #2: HSS123558; RHBDL3 #1: HSS136312; RHBDL3 #2: HSS136314; RHBDL4/RHBDD1 #1: HSS130774; RHBDL4/RHBDD1 #2: HSS130775. Negative control medium GC duplex (cat no: 462001) was purchased from Invitrogen. For rhomboid knockdowns, including where two siRNAs were combined, a final concentration of 75 nM was used. siRNAs were delivered using Lipofectamine RNAiMAX, according to the manufacturer’s instructions (Invitrogen), and incubated for indicated time periods.

### AP-TMD shedding assay

To test rhomboid cleavage of candidate substrate transmembrane domains, 5 × 10^4^ HEK293 cells were plated in one well of a 96 well plate in the presence of 30 ng each of plasmids encoding AP-TMD and 3xHA-RHBDL constructs, pre-complexed in optiMEM (Gibco) with FuGene6 HD transfection reagent (Promega), according to manufacturers instructions. Cells were left for 24 hours to attach and express protein, and then exchanged into 200 μl optiMEM overnight. AP activity was detected in the supernatants or in cell lysates (using Triton-X100 buffer) by adding equal volumes of PNPP buffer (Thermo Scientific) followed by measurement of absorbance at 405 nm on a plate reader. The percentage of the total material shed from each well (i.e. signal from supernatant divided by total signal from lysate and supernatant) was then used to calculate release, and processed as described in the figure legends. Error bars represent standard error of the mean.

### Cytosolic calcium readouts (including barium)

Cells were loaded with Fura 2 by incubating in 1 μM Fura 2-AM in external solution (145 mM NaCl, 2.8 mM KCl, 2 mM CaCl2, 2 mM MgCl2, 10 mM Dglucose, 10 mM HEPES, pH 7.4) for 40 minutes in the dark, followed by washing and incubating in external solution for another 15 minutes for full de-esterification. Ca^2+^-free solution comprised of 145 mM NaCl, 2.8 mM KCl, 2 mM MgCl2, 10 mM D-glucose, 10 mM HEPES, 0.1 mM EGTA, was applied to cells prior to Ca^2+^ image measurement, 1μM of Thapsigargin diluted to a final volume of 10 μl by Ca^2+^-free solution was applied in around 1 min after the recording started. While the trace go to the basal levels in around 10 min after Thapsigargin treatment, 2mM Ca^2+^ or Ba^2+^ were then applied. Cells were alternately excited at 356 and 380 nm, and signals were acquired every 2 sec. Calcium signals are represented by the 356 nm/380 nm ratio (R). All the images were analyzed by using IGOR Pro software.

### T cell isolation and activation assay

T cells were isolated as previously described (Breuning and Brown, 2017). In brief, using RosetteSep™, primary CD4^+^ T cells were isolated from cones from anonymous donors with approval from the National Health Service Blood and Transplant (NHSBT) and stimulated with CD3 and CD28 mAbs on Dynabeads (Thermo Fisher) in the presence of 100 U/ml IL-2 to produce T cell blasts. Primary T cell blasts were transduced with pLKO-based shRNA expressing lentivirus and selected for puromycin resistance for at least one week. Depletion of RHBDL2 mRNA was then confirmed by RT-pPCR. For functional assays, 96-well round-bottom plates were coated with varying doses of CD3 mAb (UCHT1; eBioscience). Transduced activated primary CD4^+^ T cells (5 × 10^5^ cells in 200 μl) were added, and the mixture was incubated for 18 h at 37°C. Cells were then stained with anti-CD69-allophycocyanin (APC) (Life Technologies) and analysed for percentage positivity by flow cytometry using a FACSCalibur plate reader (BDBiosciences). Dose-response curves and EC50 values were generated with GraphPad Prism.

### qRT-PCR

Total RNA was isolated from HaCaT cells by using RNeasy micro kit (Qiagen) according to the manufacturers instructions. 2000 ng RNA was used for cDNA synthesis using a cDNA Synthesis kit (PCR Biosystems) according to the manufacturers instructions. In most cases, the cDNA was then diluted 5-fold in water, except from cDNA from primary T cells, which was undiluted. qPCR was performed using TaqMan gene expression assays (Applied Biosystems) against the stated target genes in a StepOnePlus system (Applied Biosystems). GAPDH was used as a housekeeping gene for normalisation. The Applied Biosystems TaqMan probes, all purchased through Life Technologies, were as follows: RHBDL2 (Hs00983274_m1), TNF alpha (Hs00174128_m1), IL-6 (Hs00985639_m1) and GAPDH (Hs02786624_g1).

### SDS-PAGE and western blotting

Samples were typically electrophoresed at 150V on 4-12% Bis-Tris gels (Invitrogen) until the dye front had migrated off the gel (approx. 10-15 kDa). Gels were transferred onto PVDF membranes and blocked in PBS or TBS containing Tween 20 (0.05%) and 5% milk or 1% BSA, before detection with the indicated primary antibodies and species-specific HRP-coupled secondary antibodies. Band visualisation was achieved with Enhanced Chemiluminescence (Amersham Biosciences) using X-ray film. To aid quantification of Orai1 protein, all Orai1 lysate preparations were treated with PNGase (NEB) to remove all glycosylation, according to the manufacturers instruction.

### Stim1-BirA* biotin capture assay

WT and R2 KO HaCaT cells expressing pLVX-based V5-Orai1-myc and Stim1-BirA* were plated at 1 × 10^6^ in the presence of 50 μM biotin and 100 ng/ml doxycycline (to induce their expression) for 96 hours. Prior to lysis, where stated, cells were then treated with 100 nM bafilomycin A1. Cells then underwent 3x PBS washes to remove excess biotin. Cells were then lysed in RIPA buffer (50mM Tris pH 7.4, 150 mM NaCl, 1% NP40, 0.5% Sodium Deoxycholate, pH 7.4) containing complete protease inhibitor cocktail (Roche). Lysates were pulse-sonicated in an ice-water bath for 5 mins. After pelleting at 10,000 × g, clarified supernatants were incubated with 30μl high-capacity neutravidin agarose beads overnight to capture biotinylated proteins (Thermo Scientific, catalogue number 29204). Beads were then washed 3x with ice-cold RIPA buffer and eluted with 2x SDS sample buffer with excess biotin at 95°C for 15 mins. In all cases, 50% of the bead eluate and 1% lysate was loaded onto SDS-PAGE gels.

### Immunoprecipitation

HEK293 cells transfected for 24 hours with different versions of V5-Orai1-myc and 3xHA-RHBDL2-SA were grown to ~90% confluence in 10 cm plates, on the day of IP. Cells were washed 3x with PBS and then lysed in 1 ml TX-100 lysis buffer (1% Triton X-100, 150mM NaCl, 50 mM Tris-HCl, pH 7.4) supplemented with protease inhibitor cocktail (Roche). Cell lysates were cleared by centrifugation at 10,000 × g for 10 mins at 4°C. Protein concentrations were measured by a BCA assay kit (Pierce). The lysates were then immunoprecipitated for 2-3 hours with 20 μl pre-washed HA antibody-coupled beads at 4°C on a rotor. After 4-5 washes with lysis buffer, the immunocomplexes were incubated at 65°C for 15 mins in 2× SDS sample buffer. Typically, 50% of the immunoprecipitates and 1% of lysates were resolved on SDS-PAGE gels for subsequent western blotting.

### Light microscopy

HEK293 cells transfected with indicated constructs were plated on 13mm glass coverslips in 6 well dishes. Cells were washed 1x in room temperature PBS and fixed with 4% paraformaldehyde in PBS at room temperature for 20-30 mins. Fixative was quenched with 50mM NH^4^Cl for 5 mins. Cells were permeabilised in 0.2% TX-100 in PBS for 30 mins and epitopes blocked with 1% fish-skin gelatin (Sigma) in PBS for 1 hour. Coverslips were then incubated overnight with indicated antibodies in 1% fish-skin gelatin/PBS. After 3× PBS washes, coverslips were incubated with corresponding species-specific fluorescently coupled secondary antibodies (Invitrogen) for 45mins. Cells were subsequently washed 3x with PBS and 1× with H^2^O, prior to mounting on glass slides with mounting medium containing DAPI (ProLong Gold; ThermoFisher Scientific). For GFP-NFAT experiments, the fluorescent GFP signal was acquired. Images were acquired with a laser scanning confocal microscope (Fluoview FV1000; Olympus) with a 60×1.4 NA oil objective, and processed using Fiji (Image J).

### Bioinformatics

For searches based on TMD helical instability, we used HHpred in the MPI Bioinformatics Toolkit (https://toolkit.tuebingen.mpg.de/tools/hhpred). We queried the mouse proteome using Drosophila melanogaster Spitz, using the sequence surrounding and including the transmembrane domain region (PRPMLEKASIASGAMCALVFMLFVCLAFYLRFE). Most of the top hits that had an aligned transmembrane domain in the .hhr file (available upon request) were picked for the TMD screen with mouse RHBDL2. For searches of EGF domain-containing TMD proteins, we used Uniprot (https://www.uniprot.org), selecting for the presence of both transmembrane helices in a Type-I orientation and presence of an extracellular EGF-like domain. TMD regions of the hits were uniformly determined using manual searches in TMHMM Server v.2.0 (http://www.cbs.dtu.dk/services/TMHMM/) and we included three amino acids on the cytoplasmic side, and seven amino acids on the extracellular/luminal side, according to Uniprot amino acid sequence entries (https://www.uniprot.org). RHBDL2 expression data was taken from the GTEx Portal. The Genotype-Tissue Expression (GTEx) Project was supported by the Common Fund of the Office of the Director of the National Institutes of Health, and by NCI, NHGRI, NHLBI, NIDA, NIMH, and NINDS. The data used for the analyses described in this manuscript were obtained from: the GTEx Portal on 01/07/20.

---

**Figure S1:**
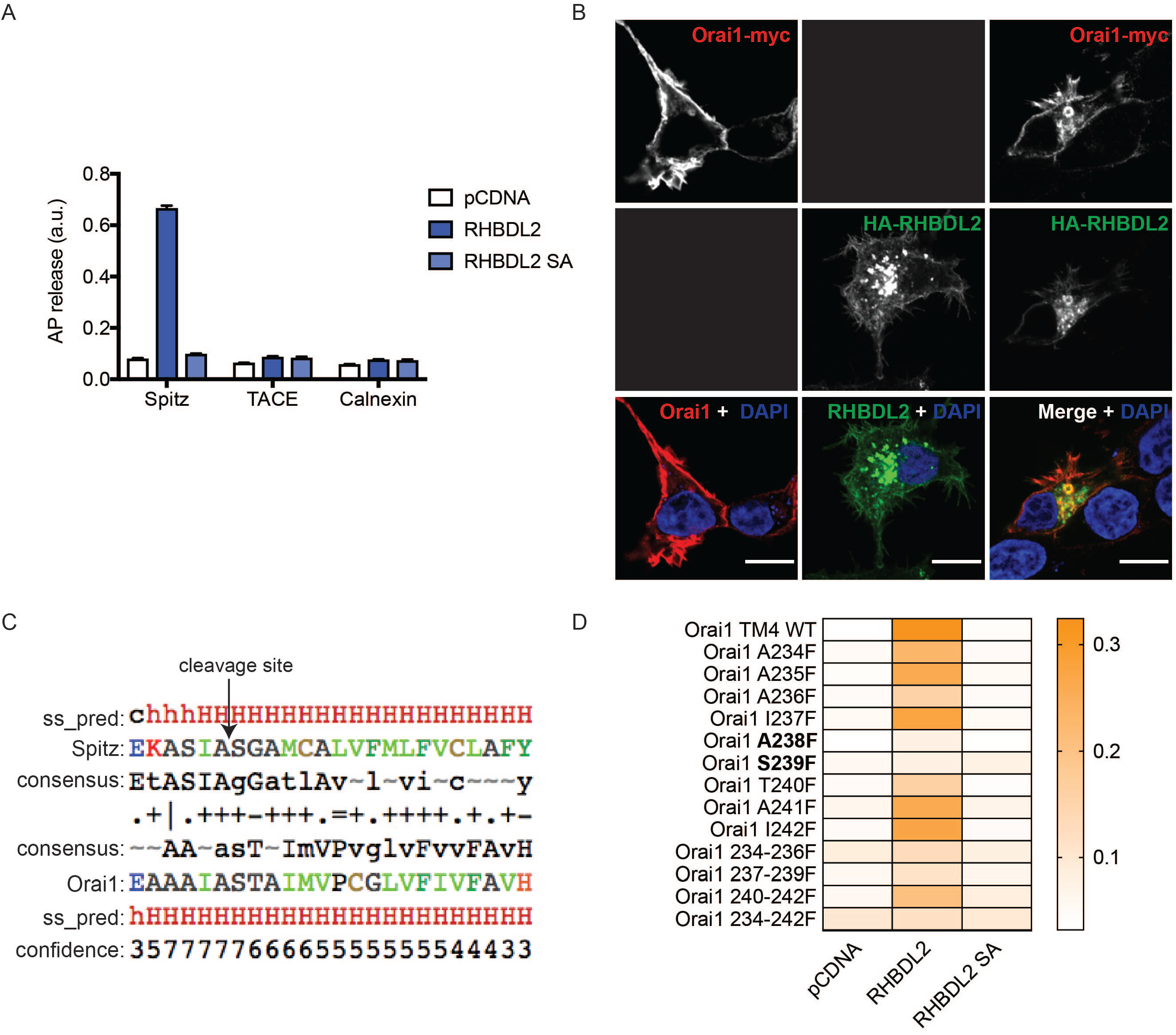
Characterisation of RHBDL2 cleavage of Orai1 TMD4. **A**. HEK293 cells were transfected with pcDNA, 3xHA-RHBDL2 or RHBDL2 SA, and indicated AP-TMDs for 48 hours. Released AP was collected over the final 16 hours. Values represent the level of released alkaline phosphatase/total alkaline phosphatase. n = 2 biological repeats for each AP-TMD. Error bars represent standard error of the mean. **B**. Immunofluorescent labelling of HA (green) and myc (red) epitopes in HEK293 cells transfected with Orai1-3xmyc and 3xHA-RHBDL2 24 hours prior to fixation. Nuclear DNA is labelled with DAPI. Black boxes indicate that this channel was not imaged. Scale bars = 10 μm. Note that internal Orai1 positive structures are only observed upon co-expression with RHBDL2. **C**. HHpred alignment of the transmembrane domain of *Drosophila* Spitz with that of TMD4 in mouse Orai1. ss_pred and confidence is the PSI-PRED secondary structure prediction and confidence values, indicating alpha-helical structure (*h = helix, c= unstructured*). The consensus line indicates the profile that was generated for the target and the query proteins. The fourth line indicates which residues align and their similarity (“I” = very good, “+” = good, “.” = neutral and “=” = clash). Residue colours: blue = acidic, red = basic, green = hydrophobic, black = polar/neutral. **D**. Cells treated as in A, but here the values have been converted into a heat-map.

**Figure S2:**
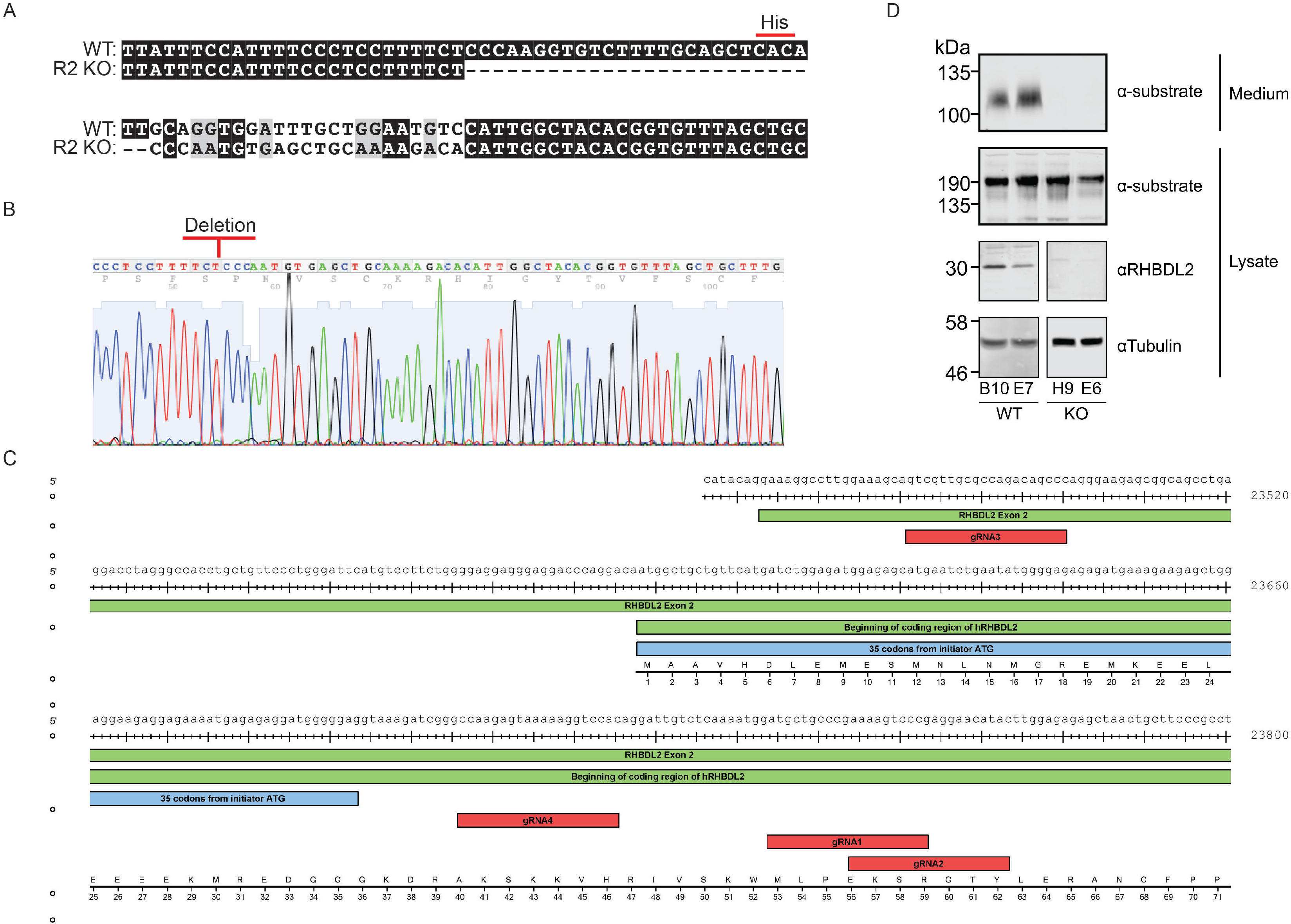
Generation of a RHBDL2 null cell lines. **A-B**. Sanger sequencing reads and alignments from WT HEK293 and R2 KO cells, showing the nucleotides flanking the catalytic histidine (CAC = His) in RHBDL2. **C.** CRISPR/Cas9 targetting scheme for RHBDL2 in HaCaT cells. **D.** Western blot for RHBDL2, and its protease activity against a confirmed shed substrate, in wild type HaCaT clones B10 and E7 and knock-out (KO) clones E6 and H9. In Figure 6C-F, only B10 and H9 clones were used.

**Figure S3:**
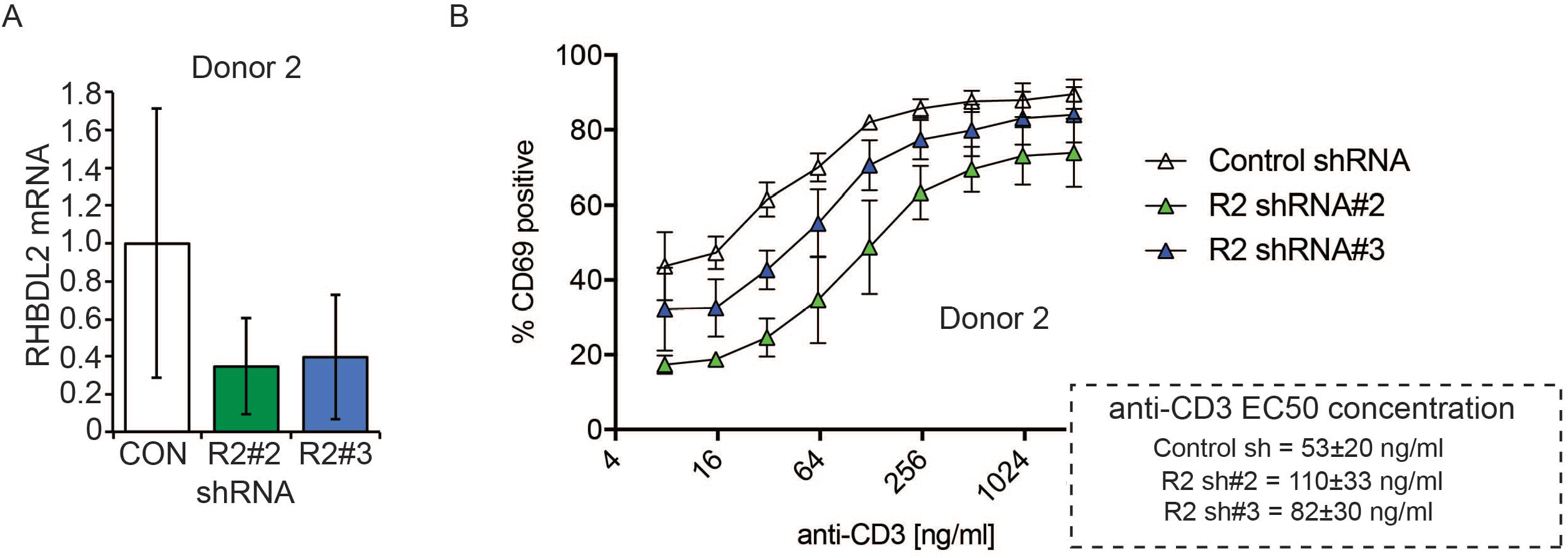
Effect of RHBDL2 depletion on primary T cell activation. **A**. TaqMan assays for RHBDL2 mRNA levels in T cells transduced with virus encoding control or RHBDL2 shRNAs. Error bars represent RQ standard error. **B.** T cell activation was measured by quantification of surface CD69 expression by FACS. CD69 expression is compared between control and RHBDL2 shRNA transduced primary CD4-positive T cells after stimulation with varying doses of platebound CD3. Each trace represents three biological replicates. In the dashed box, the calculated EC50 of anti-CD3 for each shRNA condition is indicated. Error indicates standard error of the mean.

**Figure S4:**
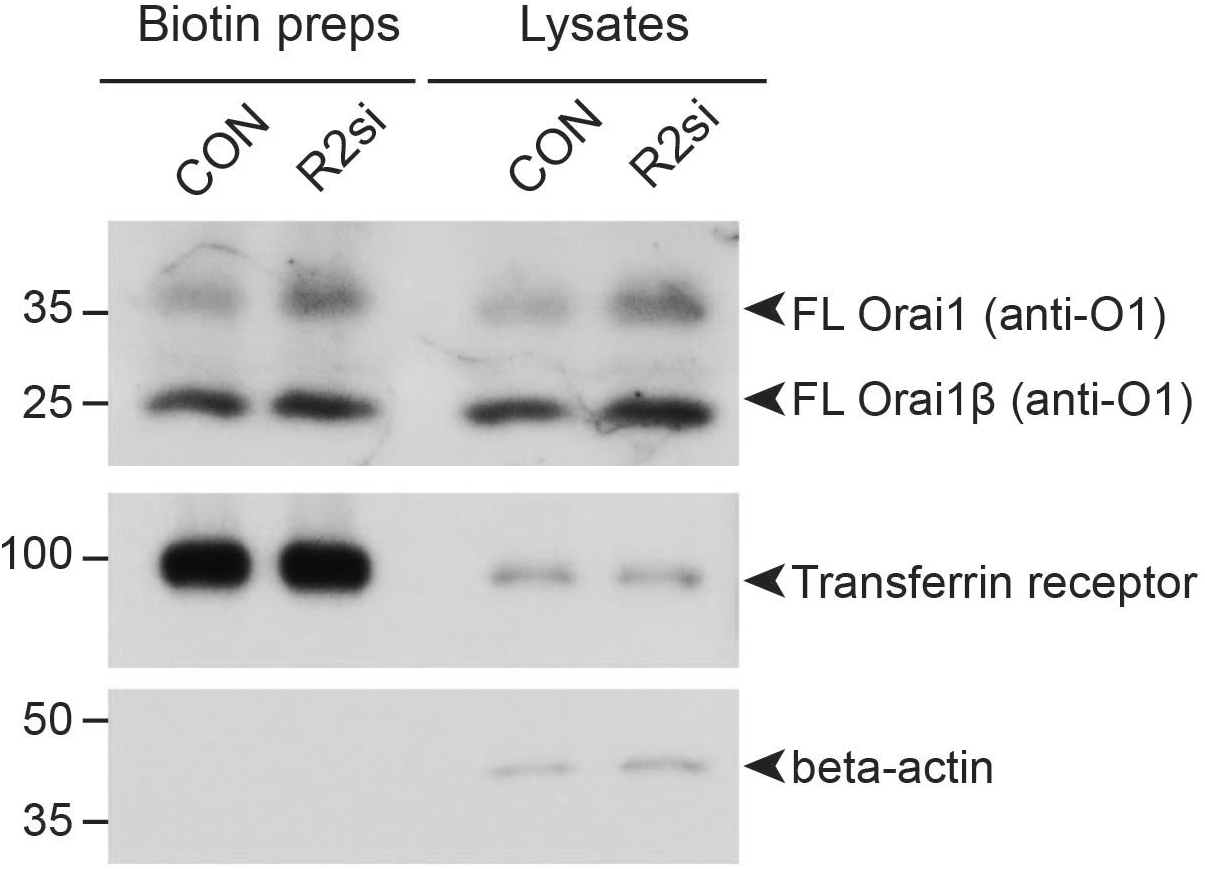
RHBDL2 controls the cell surface level of Orai1, but not transferrin receptor. Western blots of HaCaT lysates and cell surface biotinylation preps after cells were treated with control or RHBDL2 siRNAs for 72 hours, labelled for endogenous Orai1, Transferrin receptor and beta-actin. Full length Orai1 (FL) and Orai1β (FLβ) are indicated by arrowheads. Orai1β arises from alternative start sites methionine-64 or -71 (Fukushima et al., 2012).

**Table S1:**
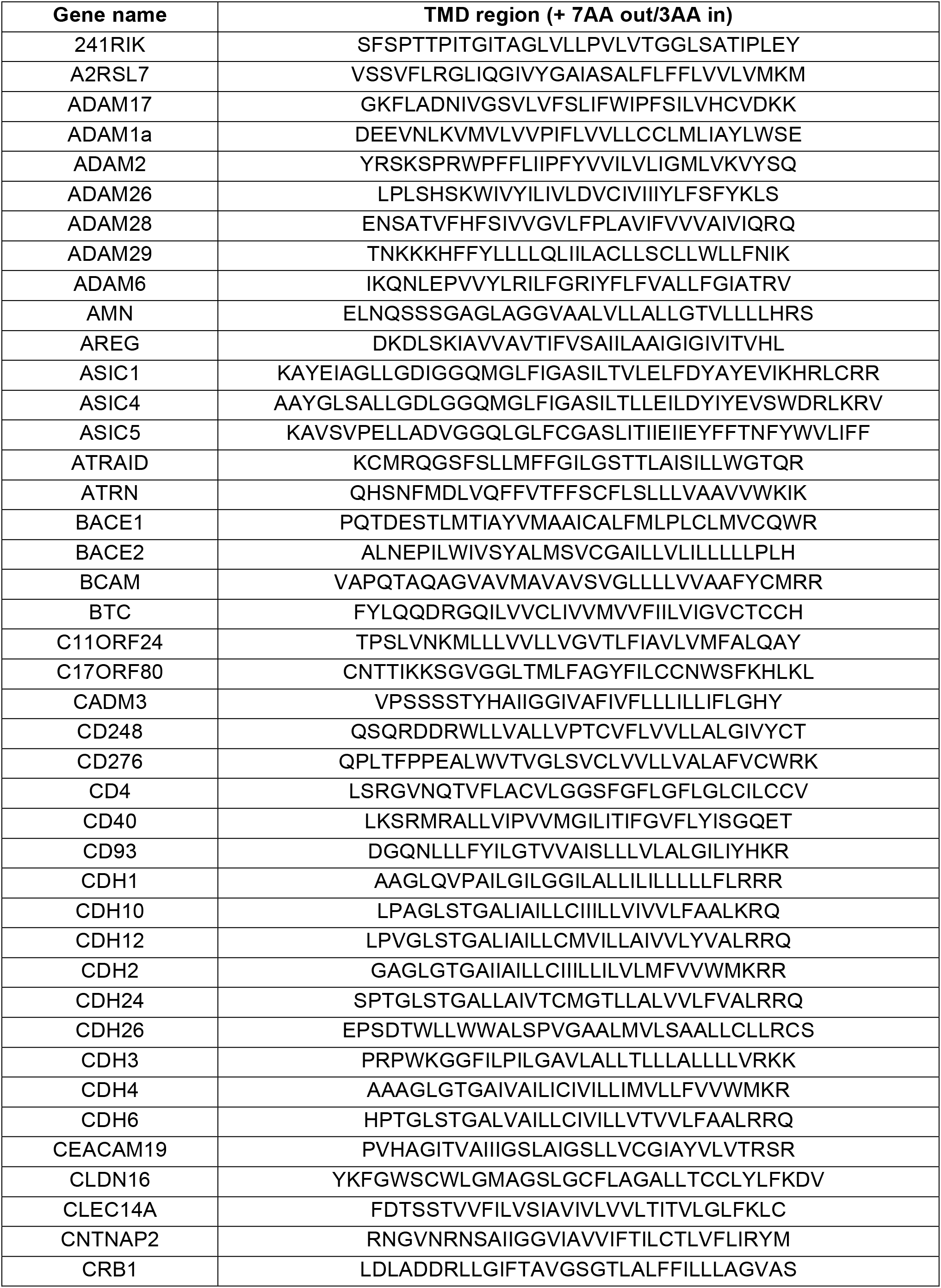
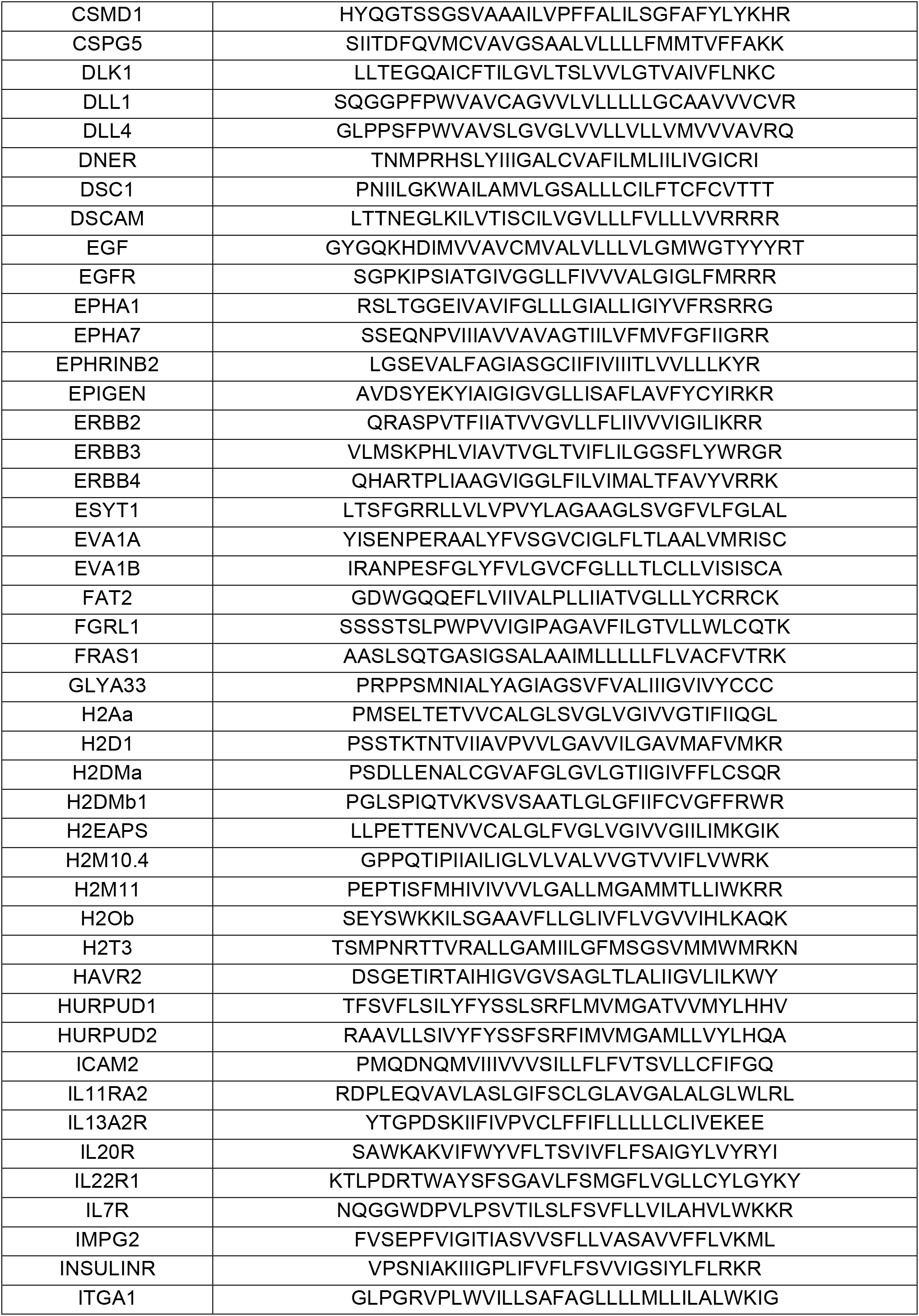
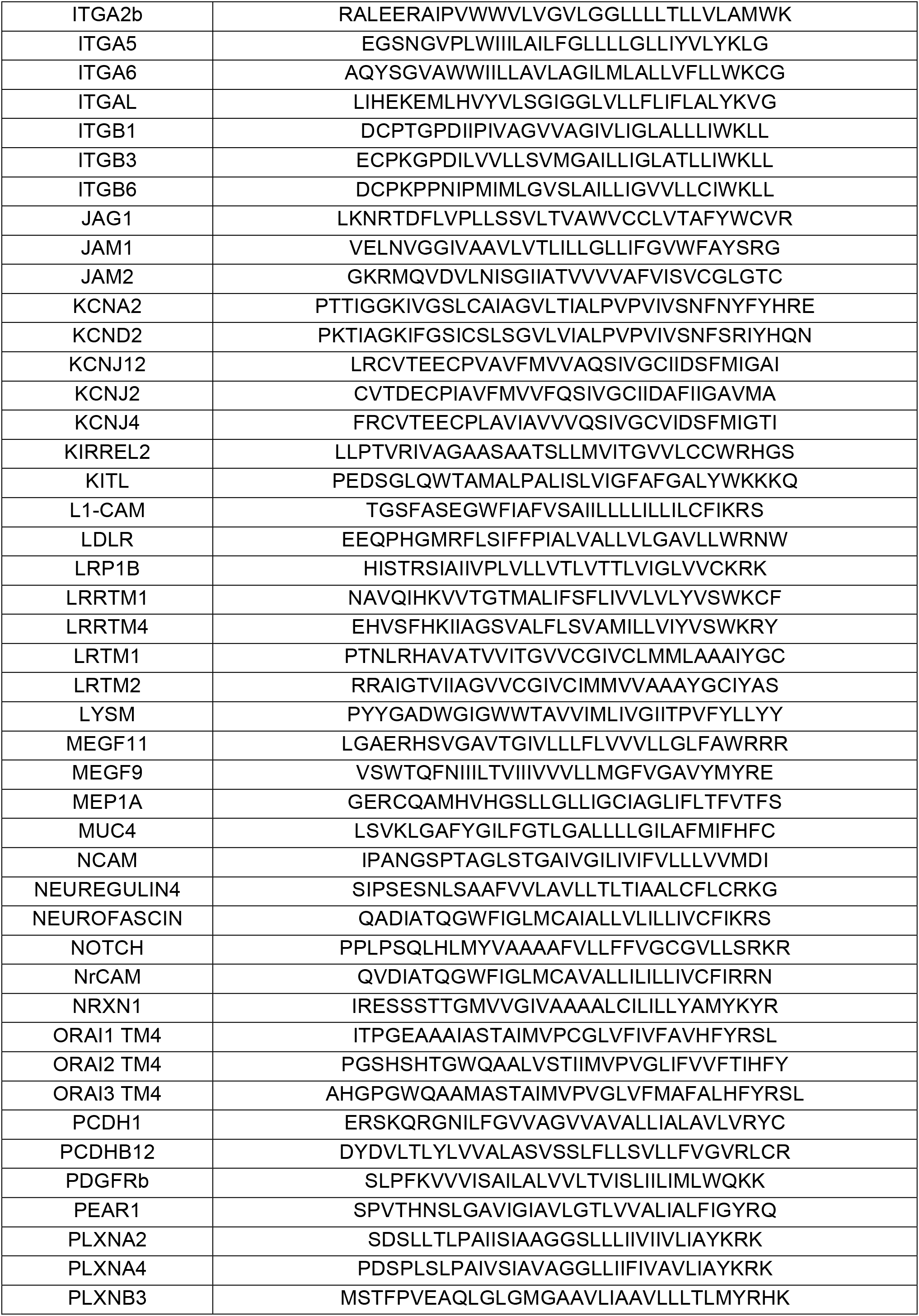
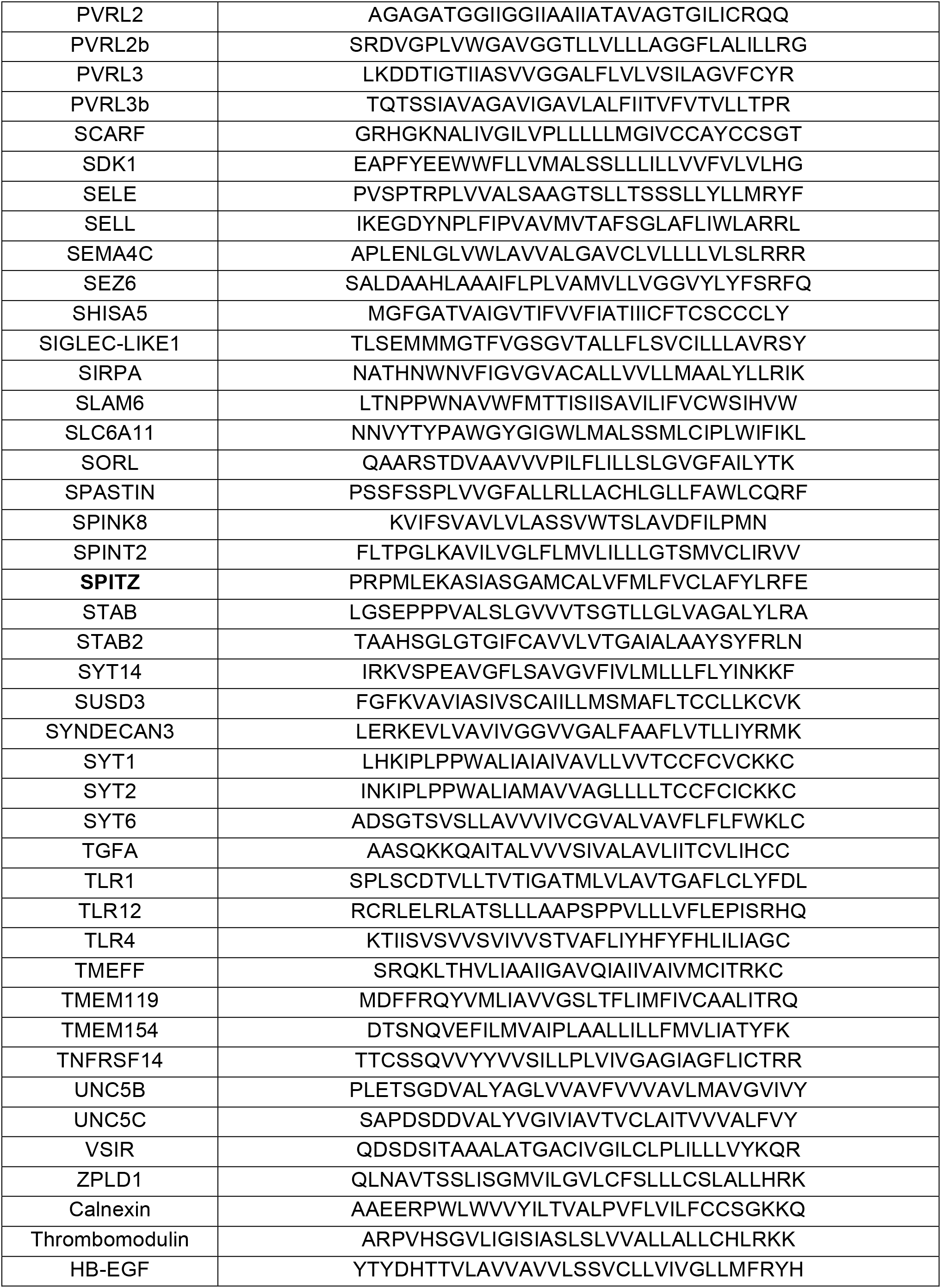
Transmembrane domain sequences used in the screen. Amino acid sequences of the TMDs and surrounding regions from indicated proteins that were used in the AP-TMD screen.

